# Plant TIR domains physically interact with EDS1 family proteins to propagate immune signalling

**DOI:** 10.1101/2023.08.02.551391

**Authors:** Jian Chen, Xiaoxiao Zhang, Maud Bernoux, John P. Rathjen, Peter N. Dodds

## Abstract

Plant Toll/interleukin-1 receptor/resistance protein (TIR) type nucleotide-binding and leucine-rich repeat immune receptors (NLRs) require Enhanced Disease Susceptibility 1 (EDS1) family proteins and the helper NLRs NRG1 and ADR1 for immune activation. TIR signalling domains possess NADase activity, producing NAM and v-cADPR from NAD^+^ *in vitro*. However, after TIR activation different small non-cyclic signalling molecules have been detected bound to EDS1/SAG101 and EDS1/PAD4 heterodimers. These molecules have not been detected in *in vitro* assays or as free molecules *in planta* and it is not clear how they are delivered to the EDS1 complexes. Here we investigate physical and functional interactions between TIR signalling domains, EDS1 family proteins and helper NLRs to clarify these signalling transduction pathways. We show that the *Nb*EDS1-*Nb*SAG101b-*Nb*NRG1 signalling pathway in *N. benthamiana* is necessary and sufficient for cell death signalling induced by six different TIR-containing NLRs from a range of plant species, suggesting this module is likely a universal requirement for TIR-NLR mediated cell death in *N. benthamiana*. We also find that TIR domains physically interact with *Nb*EDS1, *Nb*PAD4 and *Nb*SAG101 *in planta*, independently of each other. We also find evidence for direct interaction of *Nb*NRG1 with *Nb*SAG101b via its C-terminal EP domain, but not with other EDS1 family members. These data suggest a model in which physical interaction between activated TIRs and EDS1 signalling complexes facilitates efficient transfer of low abundance products of TIR catalytic activity directly to EDS1 heterocomplexes. The interaction could also alter TIR catalytic activity to favor production of the ligands recognised by EDS1/SAG101 and EDS1/PAD4.

## Introduction

The plant immune system consists of two main layers of pathogen perception (Dodds and Rathjen, 2010; Jones and Dangl, 2006; Ngou et al., 2022). Membrane-bound pattern recognition receptors (PRRs) monitor the extracellular space and can detect pathogen-derived molecules in the apoplast to trigger pattern-triggered immunity (PTI). Intracellular receptors recognise pathogen effectors that are delivered into the plant cell and activate effector triggered immunity (ETI), which is often associated with a hypersensitive response (HR) involving localised cell death. Most intracellular resistance proteins belong to the nucleotide-binding domain leucine-rich repeat (NLR) class, with either an N-terminal TIR (Toll/interleukin-1 receptor/resistance protein) or CC (coiled-coil) signalling domain (Chen et al., 2022; Qi and Innes, 2013; Zhang et al., 2017b). Cryo-EM structural analysis of the *Arabidopsis* resistance protein ZAR1 showed that the CC signalling domain adopts a four-helix bundle fold in the inactive monomer state, while effector recognition leads to formation of a pentamer in which the previously buried α1 helix of the CC domain protrudes to form the point of the funnel (Wang et al., 2019a; Wang et al., 2019b). This activated ZAR1 resistosome complex localises to the plasma membrane and has calcium-permeable cation-selective channel activity that is required for cell death signalling (Bi et al., 2021). The wheat Sr35 CC-NLR resistosome revealed a similar pentameric structure and calcium channel activity (Forderer et al., 2022), suggesting conserved activation and signalling patterns across CC-NLRs. TIR domains require self-association for cell death signalling activity (Bernoux et al 2011, Williams et al 2014, Zhang et al 2017) and exhibit an NADase catalytic activity which can cleave NAD^+^ to produce nicotinamide and a variant cyclic-ADPR (v-cADPR) *in vitro* (Horsefield et al., 2019; Wan et al., 2019). TIR signalling can be activated by forced oligomerisation through fusion to the tandem SAM domains of the human SARM1 protein, which form an octameric ring assembly (Horsefield et al., 2019), or to the mammalian NLRC4 immune receptor which forms an oligomeric inflammasome in cooperation with NAIP NLRs and a corresponding ligand (Duxbury et al., 2020; Kofoed and Vance, 2011; Zhao et al., 2011), or to the flax rust effector AvrM, which forms a stable dimer *in planta* (Bernoux et al., 2023). Activated full length TIR-NLRs RPP1 and Roq1 form tetramers where the TIR domains self-associate resulting in conformational changes that expose the NADase catalytic sites in two of the four TIR subunits (Ma et al., 2020; Martin et al., 2020), providing an explanation for how effector-driven TIR-NLR oligomerisation leads to signalling activation via its catalytic function.

TIR-NLR signalling requires two layers of downstream partners. The first layer includes the EDS1 (Enhanced Disease Susceptibility 1) family lipase-like proteins. EDS1 forms distinct heterodimers with the related protein family members PAD4 (Phytoalexin Deficient 4) or SAG101 (Senescence-Associated Gene 101) (Feys et al., 2005; Garcia et al., 2010; Wagner et al., 2013; Wiermer et al., 2005). The second layer includes two families of helper NLRs of the RPW8-like CC-NLR class, NRG1 and ADR1. These work cooperatively with either the EDS1-SAG101 or EDS1-PAD4 heterocomplexes respectively, to mediate immunity (Castel et al., 2019; Collier et al., 2011; Jubic et al., 2019; Lapin et al., 2019; Qi et al., 2018; Wu et al., 2019). EDS1, PAD4 and SAG101 share similar structures with N-terminal lipase-like domains (NLP) and C-terminal EDS1-PAD4 domains (CEP) (Feys et al., 2005; Rietz et al., 2011; Wagner et al., 2013). The heterodimer interaction is mainly mediated through a convex-concave interface formed by the protruding hydrophobic α-H helix of the EDS1 NLP domain that fits into a hydrophobic pocket of the NLP domains of SAG101 or PAD4 (Bhandari et al., 2019; Gantner et al., 2019; Wagner et al., 2013). Mutations of hydrophobic residues in the EDS1 α-H helix or in the SAG101 or PAD4 hydrophobic pockets greatly weaken the heterodimer interactions and also abrogate the signalling functions of Arabidopsis and tomato EDS1-family proteins (Gantner et al., 2019; Wagner et al., 2013).

*In vitro* NADase activity assays showed that plant TIR domains produce only nicotinamide (NAM) and a variant form of cyclic ADP ribose (v-cADPR) (Horsefield et al., 2019; Wan et al., 2019), while Yu et al (2022) showed that the L7 TIR can also hydrolyse dsDNA/dsRNA to produce 2’,3’-cAMP/cGMP. However, characterisation of Arabidopsis EDS1 heterodimers after RPP1 activation detected 2L-(5LL-phosphoribosyl)-5L-adenosine mono- or di-phosphate (pRib-AMP/ADP) and ADP-ribosylated ATP or ADP-ribosylated ADP ribose (ADPr-ATP/di-ADPR) specifically bound to pockets formed by the CEP domains of EDS1-PAD4 and EDS1-SAG101 heterodimers respectively (Huang et al., 2022; Jia et al., 2022). These TIR-catalysed small molecules allosterically promote interaction between EDS1 heterodimers and their respective NRG1 or ADR1 helper proteins. Similar to ZAR1 and Sr35, activated NRG1 and ADR1 oligomerise, associate with membranes and form calcium-permeable cation channels necessary to trigger immunity and cell death (Feehan et al., 2023; Jacob et al., 2021). The discrepancies between the *in vitro* activities of plant TIRs and the signalling molecules associated with EDS1 heterodimers are not understood and there are important questions to resolve about the relative half-lives of the different products, their cellular locations and how they are transferred between TIR-NLRs and EDS heterodimers (Locci et al., 2023).

In *N. benthamiana*, induction of cell death by the autoactive isolated *At*RPS4TIR and *At*DM2hTIR domains or by effector activation of the intact N and Roq1 TIR-NLRs requires *Nb*SAG101b but not *Nb*SAG101a or *Nb*PAD4 (Gantner et al., 2019; Lapin et al., 2019; Qi et al., 2018). *Nb*NRG1 is also required for cell death mediated by Roq1, N and *At*RPP1 in *N. benthamiana*. Here we show that the *Nb*EDS1-*Nb*SAG101b-*Nb*NRG1 signalling module in *N. benthamiana* is necessary and sufficient for cell death signalling by six additional TIR-containing NLRs from a range of plant species, suggesting it is likely a universal requirement for TIR-NLR-mediated cell death in this species. We also find that TIR domains can physically interact with the NLP domains of *Nb*EDS1, *Nb*PAD4 and *Nb*SAG101, while *Nb*NRG1 can interact with *Nb*SAG101b via its CEP domain, but this is inhibited by *Nb*EDS1/*Nb*SAG101b interaction. These data suggest that the TIR NADase-derived signal may be directly transferred to the EDS1 heterocomplex signalling modules to activate downstream signalling via exposure of a helper NLR binding surface.

## Results

### *Nb*EDS1, *Nb*SAG101b and *Nb*NRG1 are required for cell death mediated by diverse TIR-NLRs in *N. benthamiana*

We tested the ability of the TIR domains of six TIR-NLRs from flax (L6, M and M1), grapevine (Run1) and Arabidopsis (SNC1 and RPS4) to induce cell death in the *N. benthamiana* single, double or triple gene knockout lines *eds1*, *eds1/pad4*, *pad4/sag101a/sag101b* (denoted *pss*) and *nrg1*. The TIR domains were tested in different contexts that induce cell death (**Supp Fig.1**), including: autoactive TIR-alone fragments; full length TIR-NLRs co-expressed with corresponding Avr proteins; full length autoactive mutant TIR-NLRs; and TIR domain fusions to either the oligomerising SAM domain of SARM1 (Horsefield et al., 2019) or to the animal receptor NLRC4 co-expressed with NAIP5 and ligand FlaA (Duxbury et al., 2020). Individual expression of all 16 constructs induced visible cell death in wildtype (WT) *N. benthamiana* leaves, but none caused detectable cell death in *eds1* plants (**Figure 1A, Supp Fig 2**) except for those containing the RPS4 TIR domain which induced very weak cell death. The cell death induced by expression of each of these 16 constructs in *eds1* plants was restored to the level of WT leaves by the co-expression of *Nb*EDS1. The residual cell death observed for the RPS4TIR series in *eds1* plants was abolished when the NADase catalytic site glutamate was mutated to alanine (E88A) and in the case of SAM-RPS4TIR-3Myc when the wildtype SAM domain was replaced with the non-oligomerising SAM5M fusion (**Supp Fig 3**). Thus, RPS4TIR residual cell death was dependent on both oligomerisation and NADase activity.

**Figure 1.**
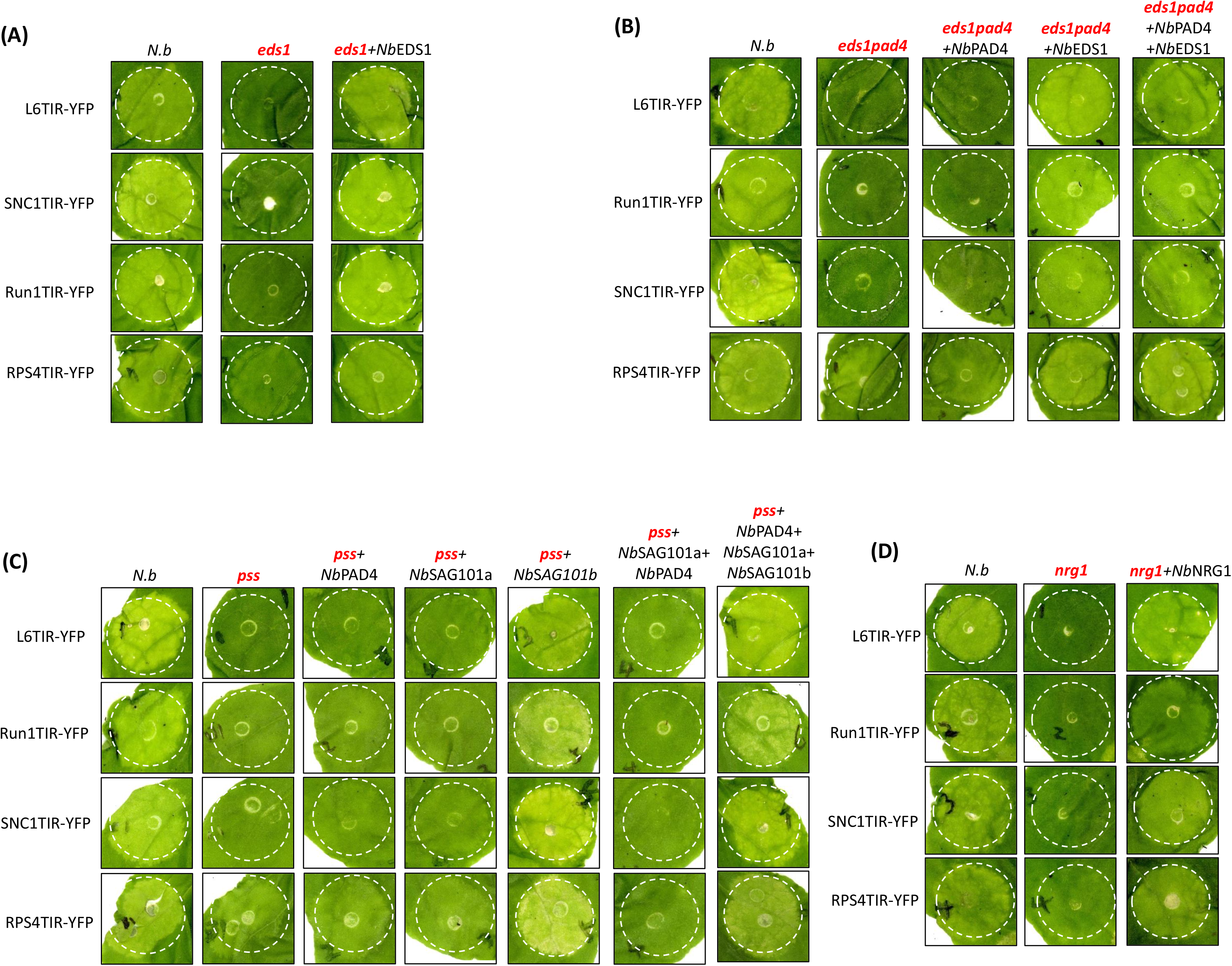
*Nb*EDS1, *Nb*SAG101b and *Nb*NRG1 are required for TIR mediated cell death in *N. benthamiana.* Complementation of TIR-mediated cell death in the *eds1* (A)*, eds1pad4* (B)*, pad4sag101a/sag101b* (*pss*) (C), and *nrg1* (D) mutant lines of *N. benthamiana*. The indicated TIR proteins fused to YFP were expressed alone in wild-type (*N.b*.) or mutant lines or in combination with *Nb*EDS1 family proteins or NRG1 fused to 3xHA in the mutant lines by Agrobacterium-mediated transient expression with the bacterial concentration at OD600 =0.5. Photos were taken at 5 days post infiltration (dpi). This experiment was repeated at least 5 times.

TIR-mediated cell death was also abolished in *eds1/pad4* double knockout plants and could be recovered with co-expression of *Nb*EDS1 but not *Nb*PAD4 (**Figure 1B, Supp Fig. 4**). Co-expression with both *Nb*PAD4 and *Nb*EDS1 complemented cell death to similar levels as *Nb*EDS1 alone. Similarly, TIR-induced cell death was abolished in leaves of the *pss* triple gene knockout and could be recovered by co-expression of *Nb*SAG101b, but not *Nb*PAD4 or *Nb*SAG101a alone or in combination (**Figure 1C, Supp Fig. 5**). Co-expression of *Nb*PAD4 and *Nb*SAG101a with *Nb*SAG101b did not affect complementation of cell death. None of the TIR-containing constructs induced cell death in *nrg1* leaves, but this phenotype was recovered by co-expression with *Nb*NRG1, although not fully to the wild-type levels (**Figure 1D, Supp Fig. 6A**). While TIR-mediated cell death could be complemented by co-expression of *Nb*NRG1 in *nrg1* plants, the cell death in WT plants was inhibited when *Nb*NRG1 was co-expressed (**Supp Fig. 6B**), which may explain why complementation was not complete. The RPS4TIR series also retained a weak cell death activity in *eds1/pad4*, *pss* and *nrg1* plants, possibly indicating a limited capacity of the RPS4 TIR to induce a weak cell death response by another unknown pathway. Overall, these data extend the observation that TIR-NLR-mediated cell death in *N. benthamiana* generally requires *Nb*EDS1, *Nb*SAG101b and *Nb*NRG1, but not *Nb*PAD4 or *Nb*SAG101a as reported previously for a few TIR-NLRs (Gantner et al 2019, Lapin et al. 2019).

Although NRG1 overexpression was previously reported to cause cell death in *N. benthamiana* (Collier et al., 2011; Peart et al., 2005; Qi et al., 2018), we did not observe this with *Nb*NRG1-3xHA or *Nb*NRG1-YFP expression in the above experiments using the pAM-PAT-35S vector. However, expression of *Nb*NRG1-3xHA from the pBIN19-35S vector caused a strong cell death phenotype (**Supp Figure 7**). Similarly, *Nb*ADR1-3xHA fusions showed strong cell death when expressed from pBIN19-35S but not from pAM-PAT-35S. In both cases this correlated with higher accumulation of protein after expression from the pBIN19-35S vector. This is consistent with the results of Qi et al., (2018) who found that lower expression of *Nb*NRG1 from its native promoter could complement TIR-NLR induced cell death in the *nrg1* mutant without autoactive cell death. The higher expressing pBIN19-*Nb*NRG1 and pBIN19-*Nb*ADR1 constructs also induced strong cell death in the *eds1*, *eds1pad4*, *pss*, and *nrg1* mutant plants (**Supp Figure 7**), consistent with the role of these helper NLRs downstream of EDS1 family proteins in *N. benthamiana.* Co-expression of the wheat CC-NLR protein Sr50 with AvrSr50, or expression of the autoactive CC domains of Sr50 or *Nb*NRG1 caused strong cell death in WT and all mutant *N. benthamiana* lines further validating that EDS1, SAG101, NRG1 are required for TIR but not CC-mediated signaling (**Supp Figure 8**).

### TIR domains associate with *Nb*EDS1, *Nb*PAD4 and *Nb*SAG101b in co-immunoprecipitation assays

To test whether signal transduction from TIR-NLRs to the EDS1 module might involve physical association of the proteins, we co-expressed TIR domain and EDS1 family proteins with either YFP or 3xHA tags and pulled down the YFP-labelled protein with anti-GFP beads to test for co-immunoprecipitation (CoIP). All four tested autoactive plant TIR-3xHA fusions co-precipitated with *Nb*EDS1-YFP, and conversely *Nb*EDS1-3xHA could be co-precipitated with each of the TIR-YFP fusions (**Figure 2A**). In contrast, neither YFP alone nor YFP fused to the CC domain of Sr50 (Sr50CC-YFP) showed interaction with *Nb*EDS1-3xHA (**Figure 2A, B**), and neither 3xHA nor YFP-tagged TIRs showed interaction with YFP alone or the Sr50CC-3xHA respectively (**Supp Figure 9A**). These TIR-*Nb*EDS1 interactions were also observed in *pss* plants **(Figure 2B** and **Supp Figure 9B**), indicating they were independent of *Nb*PAD4, *Nb*SAG101a and *Nb*SAG101b. Co-expression of a Myc-tagged *Nb*SAG101b protein did not affect the interaction between these TIR domains and *Nb*EDS1 (**Supp Figure 9C**).

**Figure 2.**
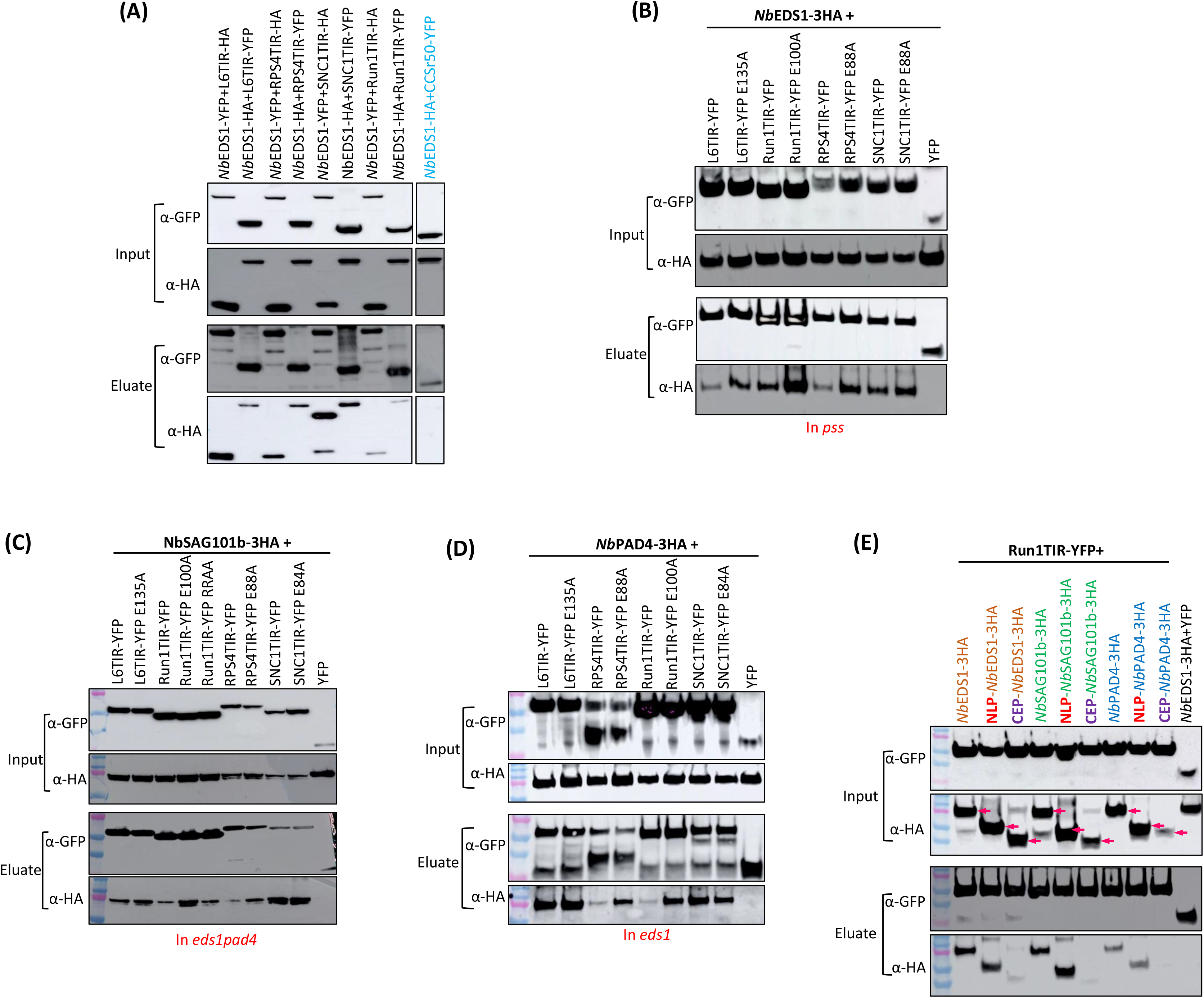
Plant TIRs interact with EDS1 family proteins in co-immunoprecipitation experiments *in planta*. *Nb*EDS1, *Nb*SAG10, *Nb*PAD4 and TIR proteins fused to YFP or 3xHA tags were transiently co-expressed in wildtype or mutant *N. benthamiana* leaves in the indicated combinations and proteins were extracted after 24 hours. Tagged proteins were detected in the extract (input) and after immunoprecipitation with anti-GFP beads (Eluate) by immunoblotting with anti-HA (α-HA) or anti-GFP (α-GFP) antibodies. YFP alone or an Sr50 CC domain YFP fusion (CCSr50-YFP) were included as negative controls as indicated. (A). Reciprocal CoIP experiments between TIR-YFP protein fusions with *Nb*EDS1-3xHA and *Nb*EDS1-YFP with TIR-3xHA expressed in wildtype plants (B). CoIP experiments of *Nb*EDS1-3xHA with TIR-YFP and NADase catalytic site mutants (E135A etc) expressed in *pad4/sag101a/sag101b* plants. (C). CoIP experiments of *Nb*SAG101b-3xHA with wildtype and catalytic mutant TIR-YFP fusions expressed in *eds1pad4* plants. (D). CoIP experiments of *Nb*PAD4-3xHA and wildtype and catalytic mutant TIR-YFP fusions expressed transiently in *eds1* mutant plants. (E). CoIP experiments of truncated *Nb*EDS1, *Nb*PAD4 and *Nb*SAG101 proteins fused to a 3xHA tag with Run1TIR-YFP expressed in *pss* (for EDS fusions) or *eds1* (for SAG101 and PAD4) plants. NLP, N-terminal lipase-like domain. CEP, C-terminal EP domain.

Mutation of the key NADase catalytic site glutamate to alanine (E to A) in the four TIR domains consistently resulted in enhanced pull-down of *Nb*EDS1 compared to the wildtype TIRs (**Figure 2B**) and was most apparent for Run1TIR (**Supp Figure 9D**). On the other hand, the Run1TIR RRAA (R64A+R65A) mutant, which has enhanced NADase activity (Horsefield et al., 2019), showed similar interaction with *Nb*EDS1 compared with Run1TIR. Mutations that disrupted the self-association of L6TIR (Bernoux et al 2011, Zhang et al 2017) did not affect the ability of L6TIR to pull down *Nb*EDS1 by CoIP (**Supp Figure 9E**). These data suggest that the TIR-EDS1 interactions do not require TIR oligomerisation or NADase activity.

Both *Nb*PAD4-HA and *Nb*SAG101b-HA co-immunoprecipitated with the TIR-YFP fusion proteins in anti-GFP pull downs in *eds1/pad4* or *eds1* plants (**Figure 2C, D**), indicating that their interactions were independent of *Nb*EDS1. Again, RPS4TIR and Run1TIR proteins containing NADase catalytic site mutations also showed stronger interaction with *Nb*PAD4 and *Nb*SAG101b, although this was not apparent for L6 and SNC1 TIRs. The plant TIRs could also interact with Arabidopsis EDS1, PAD4 and SAG101 (**Suppl Figure 10**), indicating that the interaction mode between TIRs and EDS1 family proteins might be conserved between diverse plant species. Co-IP experiments with truncated *Nb*EDS1, *Nb*PAD4 and *Nb*SAG101b fragments showed that Run1TIR interacts strongly with the NLP domain of each protein, but only weakly or not at all with the CEP domains (**Figure 2E**).

### TIR domains associate with *Nb*EDS1, *Nb*PAD4 and *Nb*SAG101 in split-luciferase and plant two-hybrid assays

We further confirmed the TIR-EDS1 family protein interactions observed in co-IP experiments by using two independent *in planta* assays. Firstly, for split-luciferase complementation (Gehl et al., 2011), the N- and C-terminal fragments of the firefly luciferase protein (nLUC and cLUC) were separately fused to L6TIR, Run1TIR and *Nb*EDS1 family proteins. All fusion proteins retained their normal function when expressed in *N. benthamiana* (autoactivity of TIR domains, **Supp Figure 11A**; and complementation of signalling mutants by EDS1 family proteins, (**Supp Figure 11B**) and were detected by immunoblotting with anti-luciferase (**Supp Figure 11C**).

To test for physical association, Run1TIR and L6TIR fused to nLUC were co-expressed with EDS1 family members fused to cLUC in either *eds1* or *pss* plants to eliminate the effects of cell death (**Figure 3**). Co-expression of nLUC-EDS1 with cLUC-SAG101b resulted in strong luciferase activity. High luciferase activity was also detected upon co-expression of Run1TIR-nLUC with *Nb*EDS1-cLUC, while co-expression with *Nb*SAG101b-cLUC and *Nb*PAD4-cLUC gave lower levels of luciferase activity (**Figure 3A**, **Supp Figure 11D**). Similarly, co-expression of L6TIR-nLUC with *Nb*EDS1 family -cLUC fusions showed enhanced luciferase activity compared to the negative controls, (L6TIR-nLUC +NRG1CC-cLUC, and CCNRG1-nLUC EDS1-cLUC family), with the L6TIR-*Nb*EDS1 and L6TIR-*Nb*SAG101 interactions stronger than observed for L6TIR-*Nb*PAD4 (**Figure 3B**). Although the Run1TIR E100A mutant showed higher luciferase activity than wildtype when paired with *Nb*EDS1-cLUC, no significant difference was seen for interactions with *Nb*SAG101 or *Nb*PAD4, or in tests with the equivalent L6TIR E135A mutant. Split-luciferase assays with truncated EDS1 family proteins showed significantly higher activity for the NLP domains fused to cLUC when co-expressed with Run1TIR-nLUC than the equivalent CEP domain fused to cLUC (**Supp Figure 11C, D**). Indeed, the NLP domains of *Nb*SAG101b and *Nb*PAD4 showed stronger interaction with Run1TIR than with the full-length proteins or the CEP domain, which is consistent with CoIP data (Figure 2E).

**Figure 3.**
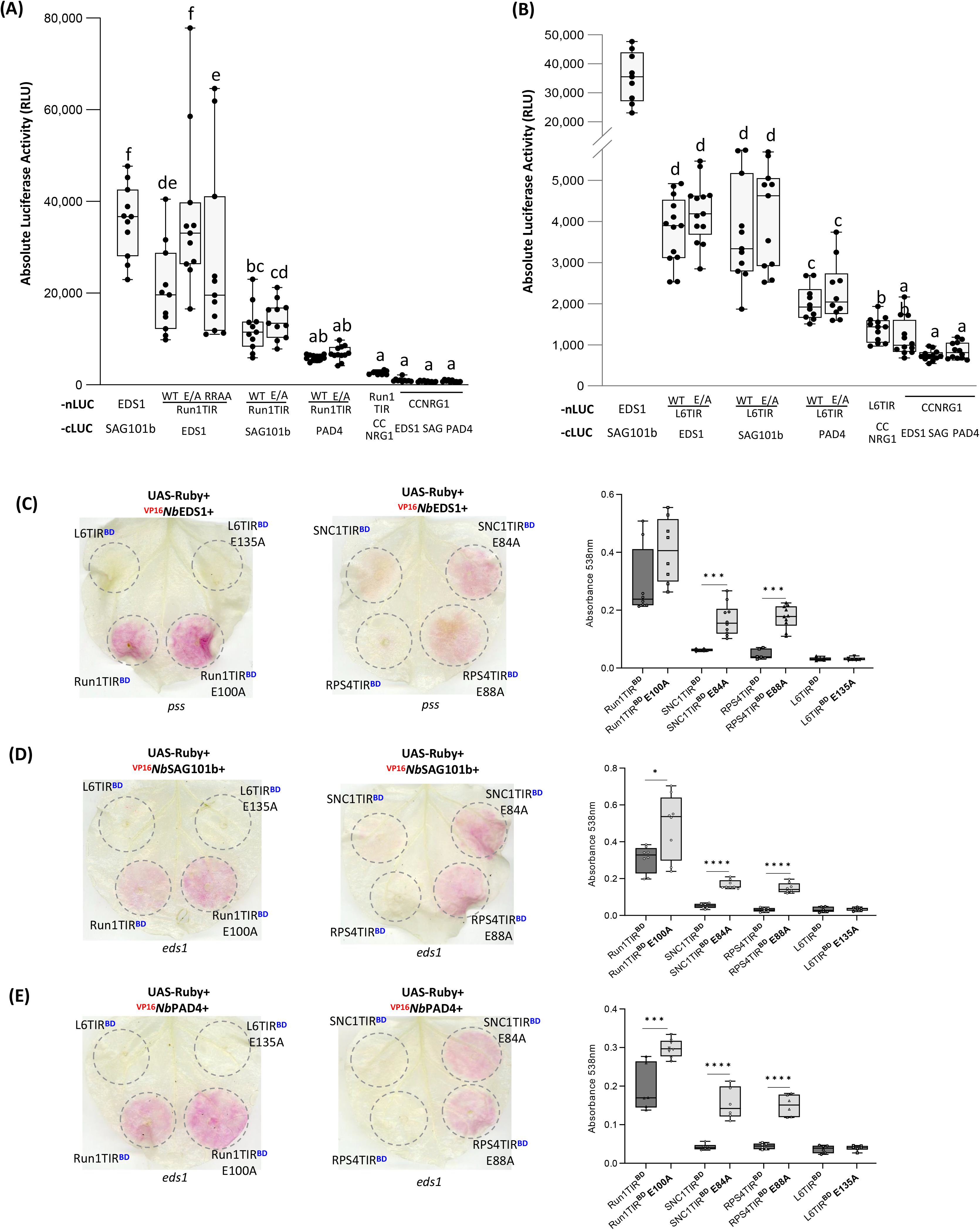
Plant TIRs interact with EDS1 family proteins in split luciferase and two-hybrid assays *in planta*. (A). Luciferase activity detected in leaf extracts expressing Run1TIR-nLUC with *Nb*EDS1, *Nb*SAG101b, or *Nb*PAD4 fused to cLUC. (B). Luciferase activity detected in leaf extracts expressing L6TIR-nLUC with *Nb*EDS1, *Nb*SAG101b, or *Nb*PAD4 fused to cLUC. EDS1-nLUC expressed with SAG101b-cLUC was used as a positive control. Co-expression of a truncated CC domain of *Nb*NRG1 (CC-NRG1) fused to -cLUC or –nLUC with the TIR-nLUC or *Nb*EDS1, SAG101 or PAD4-cLUC fusions served as negative controls. Leaf samples were taken at two dpi. The letters on the top of each column indicate statistically differences between different tests (one-way-ANOVA, LSD test, p<0.05). All experiments were conducted in *eds1* plants except the combinations of TIR-nLUC and EDS1-cLUC which were performed in *pss* plants. (C-E). Plant two-hybrid assays. Betalain accumulation was observed in leaves expressing UAS-Ruby with TIRs fused to BD and *Nb*EDS1, *Nb*SAG101b, *Nb*PAD4 fused to VP16 3-5 days after infiltration. The right columns show the absorbance at 538 nm of betalain extracted from the left leaves. Asterisks above columns indicate significant difference between wildtype TIR and the NADase site mutations. *P, 0.05, *** P, 0.001, **** P, 0.0001; Student’s t-test.

We also tested associations between TIRs and EDS family proteins using a recently-established plant two-hybrid assay. This is based on the yeast GAL4 transcription factor driving expression of the reporter *Ruby* which produces a purple pigment, betalain, as a readout for protein-protein interaction (Chen et al., 2023). Interaction assays were conducted in *eds1* and *pss* plants as above to prevent cell death. Co-expression of Run1TIR or Run1TIR E100A fused to the GAL4 DNA binding domain (Run1TIR^BD^, Run1TIR^BD^ E100A) with *Nb*EDS1 fused to the VP16 activation domain (^VP16^*Nb*EDS1) led to strong betalain accumulation (**Figure 3C**), confirming physical interactions between these proteins. The NADase mutants of SNC1 and RPS4 (SNC1TIR^BD^ E84A or RPS4TIR^BD^ E88A) also showed significant betalain accumulation when co-expressed with ^VP16^*Nb*EDS1 indicating physical interactions, while betalain production was very low for the wildtype SNC1TIR^BD^ or RPS4TIR^BD^ constructs, again indicating a stronger interaction for the catalytic mutants. Similar strength interactions were observed when these TIRs were co-expressed with ^VP16^*Nb*SAG101b or ^VP16^*Nb*PAD4 (**Figure 3D, E**). Immunoblot assays showed that all TIR^BD^ constructs accumulated to similar levels as their NADase mutants *in planta* (**Supp Figure 12**). Neither L6TIR^BD^ nor L6TIR^BD^ E135A showed interaction with ^VP16^*Nb*EDS1, ^VP16^*Nb*SAG101b or ^VP16^*Nb*PAD4 in this assay, which is likely because L6TIR is excluded from the nucleus due to its N-terminal Golgi membrane anchor (Bernoux et al., 2023). Nuclear localisation of the fusion proteins is required for activation of *Ruby* expression in this assay (Chen et al 2023). Overall, both the split luciferase and plant two-hybrid assays supported the physical association of plant TIRs with individual EDS1 family members.

### TIR domains interact with the *Nb*EDS1/*Nb*SAG101 complex in yeast

The CoIP, split-luciferase and plant two-hybrid assay results above show that plant TIRs can associate with individual EDS1 family proteins *in planta*. However, yeast two-hybrid assays failed to detect interactions between any of these TIR-EDS1 family protein combinations, except for *Nb*EDS1-BD and SNC1TIR-AD (**Supp Figure 13A and B**). On the other hand, interactions between *Nb*EDS1, *Nb*SAG101b and *Nb*PAD4 and their respective NLP but not CEP domains were detected (**Supp Figure 13C**), consistent with previous reports that EDS1 heterocomplexes are determined mainly by the N-terminal domains of EDS1 family proteins (Wagner et al., 2013). Because the EDS1 family proteins function as heterodimers, we also tested for interactions of TIRs with the *Nb*EDS1/SAG101b complex in a yeast three-hybrid (Y3H) assay, in which the BD- and AD-fused bait and prey proteins are co-expressed with a third ‘free’ protein (without AD or BD fusion). Unexpectedly, co-expression of *NbEDS1*-BD with a free *NbSAG101b* or *NbPAD4* activated yeast *HIS3* reporter gene expression allowing yeast to grow on media lacking histidine (**Supp Figure 14**), and this was dependent on the interaction since it did not occur upon co-expression with free *NbSAG101b*-LLVV or *NbPAD4*-VLI mutant proteins. However, co-expression of *NbSAG101b-BD* with free *NbEDS1* did not activate the reporter gene (**Supp Figure 14**). Thus, Y3H assays were conducted using yeast cells co-transformed with TIR-AD fusions and *Nb*SAG101b-BD along with free *Nb*EDS1 (**Figure 4**). This combination led to activation of the *HIS3* reporter gene and growth of yeast on histidine selection media for all four TIR domains tested (Run1TIR, L6TIR, RPS4TIR and SNC1TIR). However, mutation of the EDS1 heterodimer interaction surfaces (*Nb*EDS1-LTVIV or *Nb*SAG101b-LLVV) abolished these three-hybrid interactions. This indicates that in yeast, these plant TIR domains preferentially interact with the *Nb*EDS1-*Nb*SAG101b heterodimer, rather than the individual proteins as observed *in planta*. Immuno-blot analysis showed all the protein fusions were expressed in yeast (**Supp Figure 15**).

**Figure 4.**
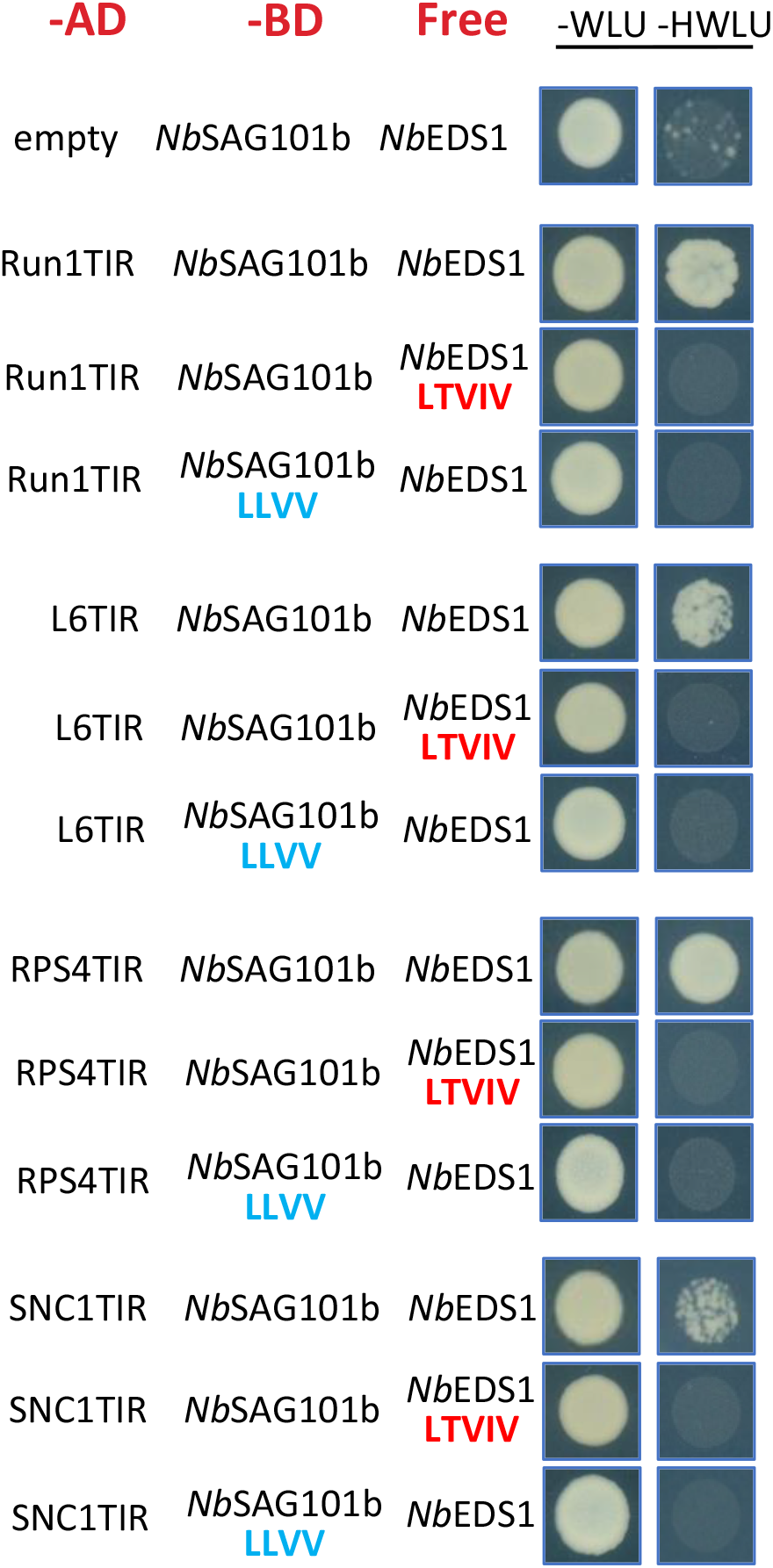
Plant TIRs interact with the *N. benthamiana* EDS1-SAG101b heterodimer in yeast. Activation of *HIS3* reporter gene expression in yeast when *Nb*SAG101b-BD plus TIRs-AD were co-expressed with free *Nb*EDS1. The selection for GAL4 activation domain (AD) constructs is leucine (L), for GAL4 binding domain (BD) constructs is -tryptophan (W) and the selection for the construct expressing free proteins in yeast is -uracil. Pictures were taken after 3 days of growth at 30lJ.

### *Nb*NRG1 associates with *Nb*SAG101b in competition with *Nb*EDS1

Because the helper NLR *Nb*NRG1 acts downstream of TIR-EDS1/SAG101b interactions, we also tested physical associations between this protein and the *Nb*EDS1 family by CoIP. *Nb*NRG1-3xHA was co-immunoprecipitated by *Nb*SAG101b-YFP, while only small amounts were detected after pull-down by *Nb*PAD4-YFP and *Nb*SAG101a-YFP, and none was detected with NbEDS1-YFP (**Figure 5A**). No interactions were detected between *Nb*ADR1 and *Nb*EDS1 family proteins under these conditions. *Nb*SAG101b could form a trimeric complex with *Nb*EDS1 and *Nb*NRG1, or two mutually exclusive *Nb*EDS1-*Nb*SAG101b and *Nb*SAG101b-*Nb*NRG1 dimeric complexes. To discriminate between these possibilities, we performed three-way CoIP assays in *eds1* mutant plants using *Nb*SAG101b-YFP as bait to pull down *Nb*NRG1-3xHA in the presence or absence of *Nb*EDS1-3Myc. As shown in **Figure 5B**, *Nb*SAG101b-YFP interacts with *Nb*NRG1 in the absence of *Nb*EDS1, but this interaction is substantially reduced when *Nb*EDS1-3Myc is co-expressed. On the other hand, co-expression of the *Nb*EDS1-3Myc-LTVIV variant, which cannot interact with *Nb*SAG101b, did not affect the interaction between *Nb*SAG101b and *Nb*NRG1. Likewise, co-expression of *Nb*EDS1-3Myc had no effect on the interaction between *Nb*NRG1 and the *Nb*SAG101b-LLVV variant which cannot interact with *Nb*EDS1. As expected, *Nb*SAG101b-YFP pulled down *Nb*EDS1-3Myc, but not *Nb*EDS1-3Myc-LTVIV, while *Nb*SAG101b-YFP-LLVV did not pull down *Nb*EDS1-3Myc. Thus, the presence of *Nb*EDS1 inhibits the interaction between *Nb*SAG101b and *Nb*NRG1 and this is dependent on its ability to form a heterocomplex with *Nb*SAG101. This suggests that the *Nb*EDS1-SAG101b and *Nb*SAG101b-NRG1 complexes are independent and mutually exclusive. In the absence of TIR-mediated signaling, the *Nb*EDS1-SAG101b interaction appears to be the favoured form as expression of *Nb*EDS1 had a strong effect on inhibiting the *Nb*SAG101b-NRG1 interaction, while expression of NRG1 had only a weak effect in reducing the EDS-SAG101b interaction.

**Figure 5.**
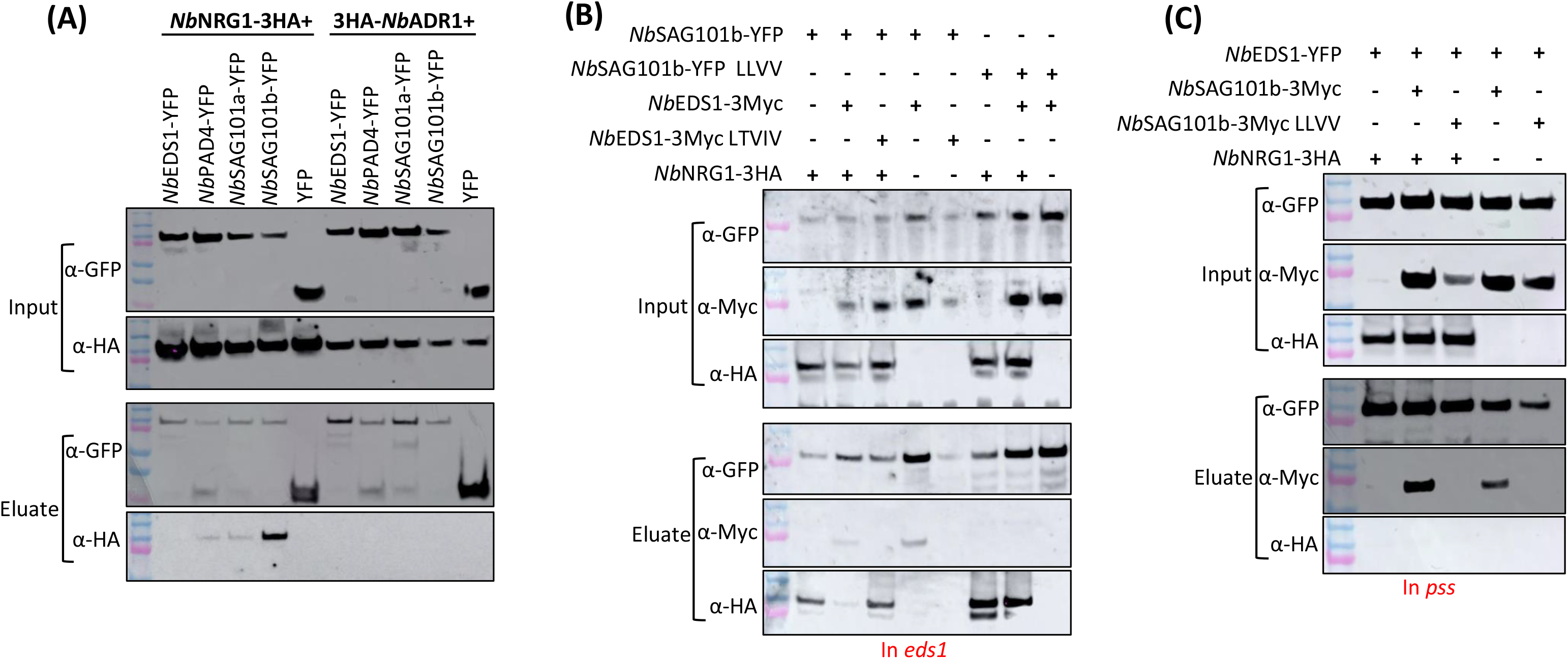
Interaction of *N. benthamiana* EDS1 family proteins with helper RNLs. (A). CoIP experiments between *Nb*EDS1-YFP, *Nb*PAD4-YFP or *Nb*SAG101-YFP with *Nb*NRG1-3xHA or *Nb*ADR1-3xHA. The indicated protein fusion combinations were expressed transiently in wildtype *N. benthamiana* plants. Tagged proteins were detected in the extract (input) and after immunoprecipitation with anti-GFP beads (Eluate) by immunoblotting with anti-HA (α-HA) and anti-GFP (α-GFP) antibodies. (B). Competitive CoIP testing interactions between *Nb*SAG101b-YFP, *Nb*SAG101b-YFP LLVV and *Nb*NRG1-3xHA, *Nb*EDS1-3Myc and *Nb*EDS1-3Myc LTVIV. Protein combinations indicated above the blot (+ construct agro-infiltrated; - non agro-infiltrated construct) were expressed in *eds1* plants. (C). Competitive CoIP for interactions between *Nb*EDS1-YFP with *Nb*NRG1-3xHA and *Nb*SAG101-3Myc and *Nb*SAG101-3Myc LLVV. The indicated protein fusion combinations were expressed transiently in *pss* plants.

As a further test for a possible trimeric complex, CoIP assays were carried out in *pss* mutant plants using *Nb*EDS1-YFP as bait to pull down *Nb*NRG1-3xHA and/or *Nb*SAG101b-3Myc. As previously, *Nb*EDS1-YFP did not pull down *Nb*NRG1-3xHA but did pull down *Nb*SAG101b-3Myc. Neither the presence of *Nb*SAG101b-3Myc or *Nb*SAG101b-3Myc LLVV variant could mediate *Nb*EDS1-YFP to pull down *Nb*NRG1-3xHA (**Figure 5C**).

### The CEP domain of *Nb*SAG101b and the NB-LRR region of *Nb*NRG1 mediate their interaction

To further characterise the *Nb*NRG1-*Nb*SAG101b interaction, we tested various subdomains of each protein for interaction. The CEP domain of *Nb*SAG101b showed stronger interaction with *Nb*NRG1 in CoIP assays than the NLP domain, albeit weaker than full length *Nb*SAG101b (**Figure 6A**), suggesting that the interaction is mediated mainly by the CEP domain. To further test this, the NLP and CEP domains of *Nb*SAG101b were swapped with the corresponding domains from *Nb*PAD4 and *Nb*SAG101a to generate four chimeric proteins: NSb:CP, NSb:CSa, NP:CSb and NSa:CSb (**Supp Figure 16**). The chimeric proteins were first tested for interaction with *Nb*EDS1. All four chimeras showed strong interactions with *Nb*EDS1 similar to the wildtype *Nb*PAD4 and *Nb*SAG101 proteins (**Figure 6B**), confirming that they can form heterocomplexes with *Nb*EDS1. However, only the wildtype *Nb*SAG101b and chimeras containing the *Nb*SAG101b CEP domain, NP:CSb and NSa:CSb, could co-immunoprecipitate *Nb*NRG1-3xHA, whereas *Nb*SAG101a, *Nb*PAD4, *Nb*EDS1 and chimeras with either the *Nb*PAD4 or *Nb*SAG101a CEP domains (NSb:CP and NSb:CSa) failed to interact with *Nb*NRG1 (**Figure 6C**). The NP:CSb and NSa:CSb chimeric proteins, which both interacted with *Nb*NRG1, also recovered cell death induction by Run1TIR-YFP in the *N. benthamiana pss* line (**Figure 6D**). Expression of the NSb:CP and NSb:CSa constructs did not support cell death signalling in this background. Thus, the CEP domain of *Nb*SAG101b determines the specificity of interaction with *Nb*NRG1 and the ability to activate TIR-induced cell death in *N. benthamiana*.

**Figure 6.**
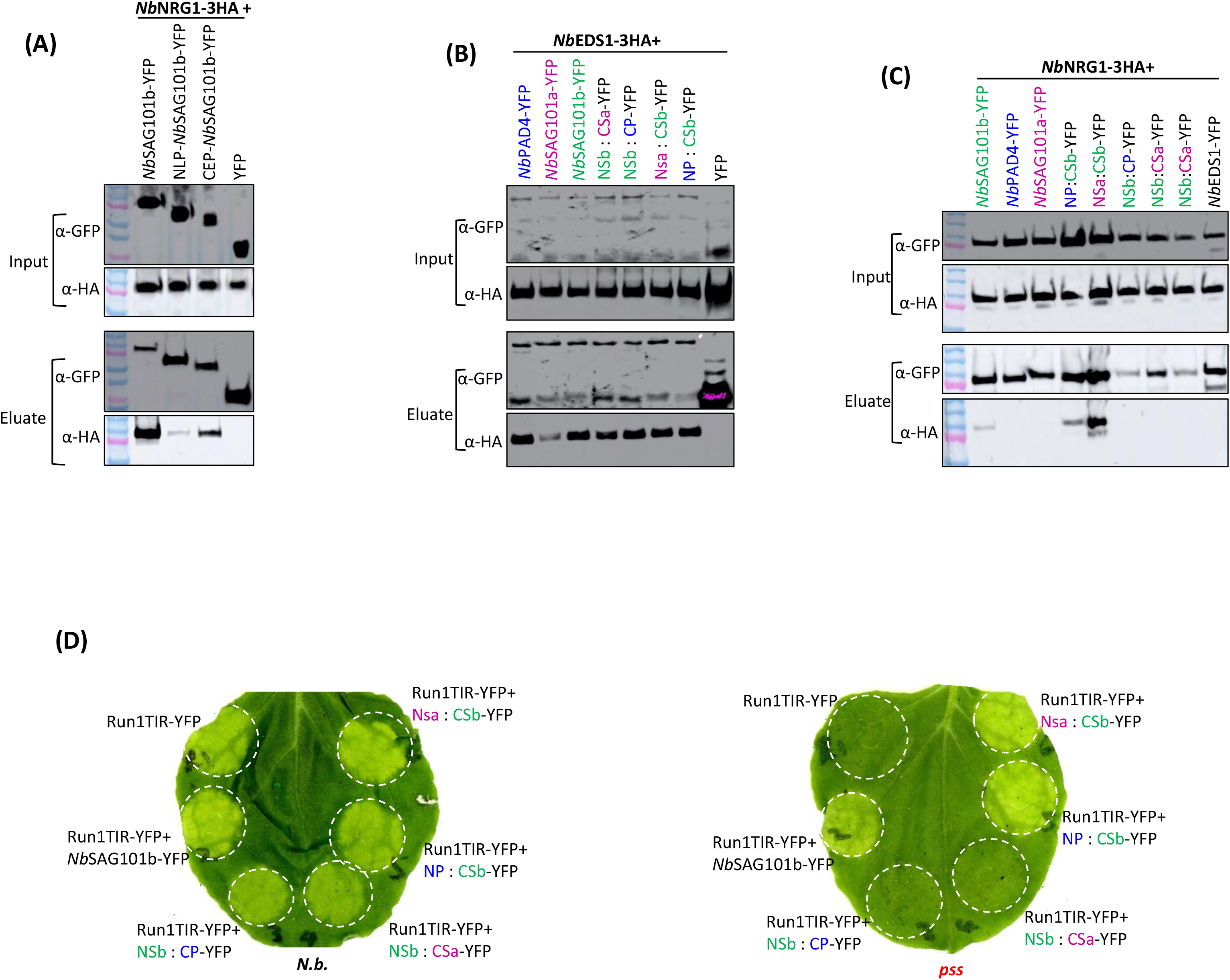
Functional analysis of EDS1-family chimeric proteins. (A). CoIP assays between *Nb*SAG101b, the N-terminal lipase-like (NLP) domain of *Nb*SAG101b and the C-terminal EP (CEP) domain of *Nb*SAG101b (fused to YFP tags) with *Nb*NRG1-3xHA. (B) and (C). CoIP assays between *Nb*EDS1-3xHA and *Nb*NRG1-3xHA with *Nb*EDS1-*Nb*SAG101b, *Nb*EDS1-*Nb*SAG101a, *Nb*EDS1-*Nb*PAD4 and the chimeric proteins. (D). Complementation of Run1TIR-YFP cell death by *Nb*SAG101b and the domain swap chimeric proteins in *pss* mutant plants. The agrobacterium concentration was OD600 = 0.5. All combinations caused similar cell death phenotypes in wildtype *N. benthamiana*.

We also tested the interaction between *Nb*SAG101b and different domains of *Nb*NRG1 (**Figure 7A**). *Nb*SAG101b-YFP could pull down full length *Nb*NRG1-3xHA as well as the NB (aa 219-609), LRR (aa 604-850) and NB-LRR (aa 219-850) fragments, but not the CC (aa 1-182) domain. Similarly, all of the NRG1 domain fragments except for the CC domain interacted with the *Nb*SAG101b chimeric proteins NSa:CSb and NP:CSb (**Figure 7B**). Surprisingly, the separated NB and LRR domains of *Nb*NRG1 each interacted with all members of *Nb*EDS1 family, including *Nb*PAD4, *Nb*SAG101a and the four chimeric proteins, although the interaction with *Nb*EDS1 was weak (**Figure 7C, D**). However, the NB-LRR fragment of NRG1 showed interaction only with *Nb*SAG101b and the NSa:CSb and NP:CSb chimeras (**Figure 7E**), similar to the full length NRG1 protein. Thus, it appears that the combination of the NB and LRR domains confers the specificity of NRG1 binding to *Nb*SAG101b.

**Figure 7.**
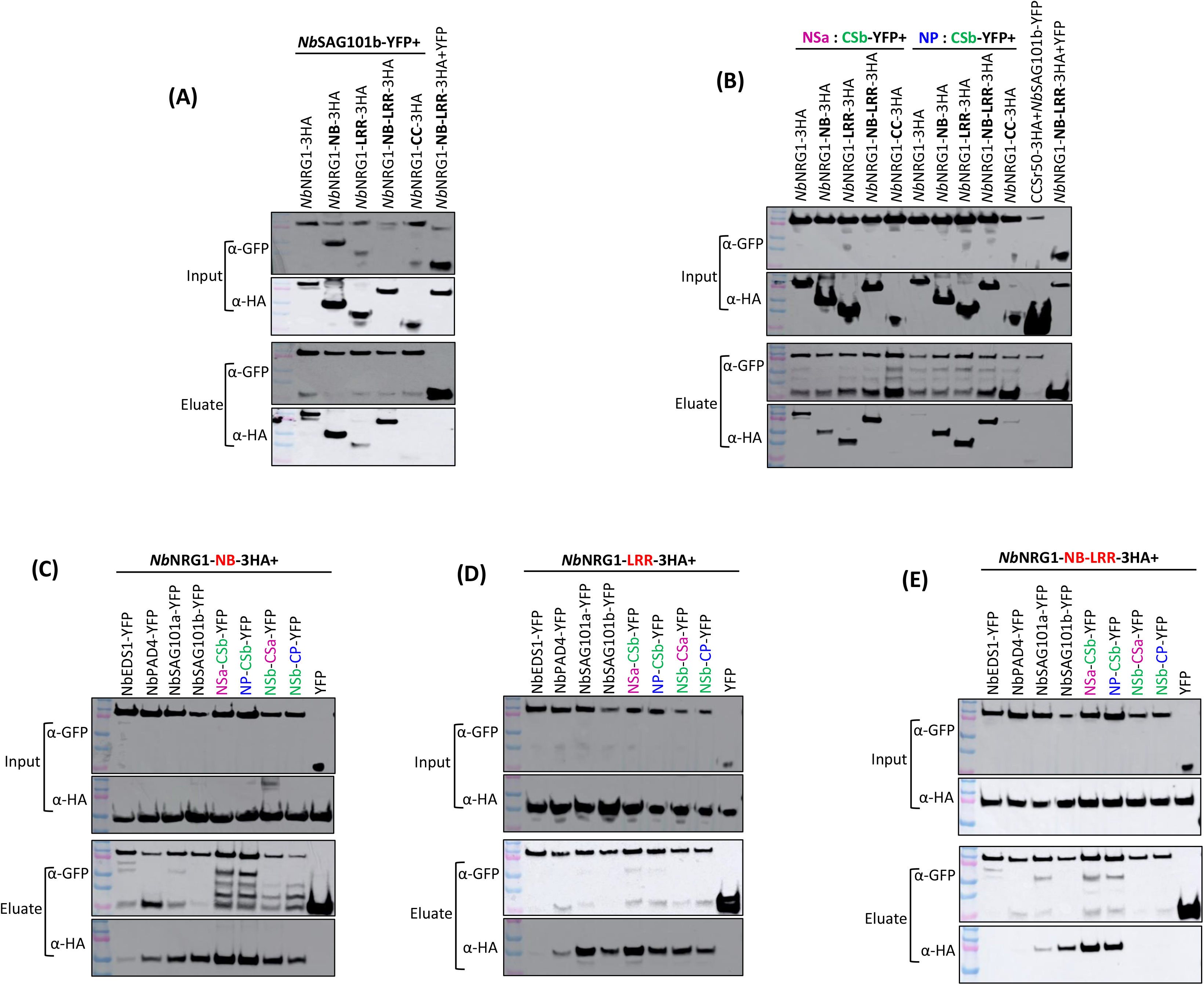
Interactions between *Nb*NRG1 domains and EDS1 family proteins. (A) and (B). CoIP assays between *Nb*NRG1 and its subdomains fused to a 3xHA tag in combination with *Nb*SAG101b chimeric proteins NSa:CSb and NP:CSb fused to a YFP tag. (C)-(E). CoIP assays of *Nb*NRG1 domains NB, LRR, and NB-LRR in combination with *Nb*EDS1 family and chimeric proteins.

## Discussion

EDS1 forms mutually exclusive heterocomplexes with PAD4 or SAG101 and acts downstream of TIR-NLR immune receptors to transduce immune signals to the downstream helper NLRs NRG1 and ADR1 (Castel et al., 2019; Rietz et al., 2011; Wagner et al., 2013; Wu et al., 2019). Some TIR-NLR receptors have been reported to require only *Nb*EDS1, *Nb*SAG101b and *Nb*NRG1 for cell death in *N. benthamiana*, but not *Nb*SAG101a or *Nb*ADR1 (Gantner et al., 2019; Lapin et al., 2019; Qi et al., 2018). Here we showed that the *Nb*EDS1-*Nb*SAG101b-*Nb*NRG1 signalling module in *N. benthamiana* is commonly required for cell death mediated by a broader set of TIR domains from diverse plant species expressed in different cell death activating contexts (**Figure 1 and Supp Figures 1-6**). In contrast, *Nb*PAD4 and *Nb*SAG101a do not contribute to cell death induction despite forming heterocomplexes with *Nb*EDS1. An exception was observed for cell death induced by the RPS4 TIR domain, which was reduced but not fully abolished in all of the mutant lines. This residual activity was dependent on TIR catalytic activity and oligomerisation, so the RPS4 TIR may partially bypass *Nb*EDS1/*Nb*SAG101/*Nb*NRG1 to signal via an unknown mechanism(s).

### TIR domains interact physically with EDS1 family proteins

The *Arabidopsis* full length TIR-NLRs RPS4, RPS6 and SNC1 have been reported to interact with EDS1 in Co-IP experiments (Bhattacharjee et al., 2011; Heidrich et al., 2011; Huh et al., 2017; Kim et al., 2012). We found that the TIR domains of L6, Run1, RPS4 and SNC1 associated with *Nb*EDS1, *Nb*PAD4 and *Nb*SAG101 in Co-IP experiments (**Figure 2**), and this was confirmed by *in planta* two-hybrid assays and also by split-LUC assays for Run1TIR and L6TIR (**Figure 3, Supp Figure 11, 12**). Similar interactions were observed between these TIR domains and the *Arabidopsis* EDS1 family proteins (**Supp Figure 10**). In these *in planta* experiments, the TIR interactions with *Nb*EDS1 were independent of *Nb*PAD4 and *Nb*SAG101, and likewise the TIR interactions with *Nb*PAD4 and *Nb*SAG101 were independent of *Nb*EDS1. However, in yeast two-hybrid assays (**Figure 4**), TIR interactions were detected only with *Nb*EDS1 and *Nb*SAG101 complexes, but not with the individual proteins. In the *in planta* CoIP assays, co-expression of *Nb*SAG101 did not affect the TIR domain interaction with *Nb*EDS1 (**Supp Figure 9C**), suggesting that the two interactions are not mutually exclusive and could occur simultaneously. This may be required for detection in the yeast assay system. Moreover, mutations that affect L6TIR oligomerisation also did not affect its interaction with *Nb*EDS1 (**Supp Figure 9C**), suggesting that both monomeric and oligomeric TIR domains could interact with EDS1 family proteins. Run1TIR interacts primarily with the NLP domain fragments of *Nb*EDS1, *Nb*PAD4, *Nb*SAG101b rather than their CEP domain fragments (**Figure 2E and Supp Figure 11D**). Given the tetrameric structure of the activated TIR domains in RPP1 and Roq1 resistosomes (Ma et al., 2020; Martin et al., 2020), and the stable heterodimers formed by EDS1 family members, it is possible that both partners of an EDS1/SAG101b heterodimer could interact directly with two different TIR domain subunits of an activated TIR-NLR resistosome (**Supp Figure 17**). While monomeric TIR domains are capable of interacting with either the EDS1 or SAG101 subunits of EDS1-SAG101 heterocomplex, only the tetramer is catalytically active and able to interact with both subunits. Interaction of EDS1 heterodimers with inactive monomeric TIR domains may facilitate their recruitment by activated resistosomes to allow substrate-mediated signalling.

The pRib-AMP/ADP and ADPr-ATP/di-ADPR products detected in TIR-activated EDS1-PAD4 and EDS1-SAG101 complexes were not detected as free molecules *in planta* (Huang et al., 2022; Jia et al., 2022), nor in *in vitro* enzyme assays (Horsefield et al., 2019; Wan et al., 2019). The observed interaction between TIR domains and EDS1 family heterodimers may help explain the discrepancy between these results in several ways. Firstly, given the instability or low abundance of the pRib-AMP/ADP and ADPr-ATP/di-ADPR products *in planta*, physical interaction may be required to allow efficient transfer of these molecular signals from activated TIR domains to EDS1 protein complexes. Secondly, the enzymatic activity of the TIR domains may be altered when in complex with EDS1 heterodimers such that these signalling molecules become the favoured products in preference to v-cADPR observed in *in vitro* assays of isolated TIR domains. Thirdly, it is possible that the activated TIR enzymes act on a substrate pre-bound to the EDS1 complexes. These hypotheses are not mutually exclusive and may all apply. Interestingly, the TIR NADase catalytic site mutations showed stronger interactions with *Nb*EDS1 family proteins than did the wildtype proteins in several assays. A possible explanation is that these mutations enhance the protein complex stability by preventing loss of a co-bound substrate or dissociation of the TIR from the complex after detection of the catalytic product by *Nb*EDS1/*Nb*SAG101b.

### NRG1 interacts with SAG101b through the CEP domain

We found that *Nb*NRG1 interacted with *Nb*SAG101b in CoIP experiments but not with *Nb*PAD4 or *Nb*SAG101a (**Figure 5 A and B**). This is consistent with the requirement for the *Nb*EDS1/*Nb*SAG101b/*Nb*NRG1 signalling module for TIR-NLR signalling in *N. benthamiana*, and the specific role of NRG1 downstream of EDS/SAG101 but not EDS1/PAD4 (Locci et al., 2023). Although *Nb*NRG1 was previously reported to interact with *Nb*EDS1 in CoIP (Qi et al., 2018), interactions with other EDS1 family members were not previously tested. However, we did not detect any interaction between *Nb*EDS1 and *Nb*NRG1. This may be explained by higher expression levels of *Nb*NRG1 in the Qi et al. (2018) study, which showed very strong autoactivity. The *Nb*EDS1/*Nb*SAG101b and *Nb*NRG1/*Nb*SAG101b interactions appeared to be mutually exclusive, since *Nb*EDS1 effectively competed with *Nb*NRG1 to form complexes with *Nb*SAG101b and this competition did not occur with the non-interacting mutants *Nb*EDS1-LTVIV or *Nb*SAG101b-LLVV (**Figure 5 C**). Thus, under normal cellular conditions *Nb*EDS1 likely sequesters *Nb*SAG101b and prevents interaction with *Nb*NRG1 in the absence of TIR activation. *Nb*NRG1 interacts preferentially with the CEP domain of *Nb*SAG101b, and domain swaps between *Nb*SAG101b and *Nb*PAD4 and the non-functional *Nb*SAG101a revealed that the CEP domain both controls this interaction specificity and complements cell death signalling (**Figure 6**). These conclusions are consistent with previous work by Gantner. et al. (2019) who tested similar domain swaps between the tomato *Sl*SAG101a and *Sl*SAG101b proteins and found that the C-terminal domain of *Sl*SAG101b was sufficient to restore activity to the non-functional *Sl*SAG101a. Similarly, chimeric proteins between Arabidopsis PAD4 and SAG101 revealed that the *At*SAG101 CEP domain is necessary for Roq1 induced cell death in *N. benthamiana* (Gantner et al., 2019; Lapin et al., 2019).

It is important to note that the interactions between *Nb*NRG1 and *Nb*SAG101b observed here occur in the absence of cell death signalling, which requires TIR domain enzymatic activity. Thus, while these interactions may define the specificity of *Nb*NRG1 interaction with *Nb*SAG101b, activation of *Nb*NRG1 requires the presence of the activated *Nb*EDS1-*Nb*SAG101b complex bound to ADPr-ATP. Indeed, binding of di-ADPR or ADPr-ATP by the *At*EDS1-*At*SAG101 heterodimer induces allosteric rotation of the *At*SAG101 CEP domain, which is proposed to trigger immunity activation by promoting interaction with *At*NRG1A (Huang et al., 2022; Jia et al., 2022). Thus, a likely scenario is that this alteration in the *Nb*EDS1/*Nb*SAG101b complex after TIR activation leads to the exposure of a binding surface of the *Nb*SAG101b CEP that interacts directly with *Nb*NRG1. This surface would be exposed in the free monomeric *Nb*SAG101b expressed in the Co-IP assays here but would not be available in the pre-activation *Nb*EDS1/*Nb*SAG101 complex present under normal cellular conditions. Structural data on ZAR1 and Sr35 resistosomes revealed that the ligand bound in the active state would sterically clash with the NB domain position in the inactive state, thereby promoting resistosome assembly (Forderer et al., 2022; Wang et al., 2019b). Monomeric *Nb*SAG101b may bind to inactive *Nb*NRG1 without causing such a steric clash, while binding of the larger *Nb*EDS1/*Nb*SAG101 complex would promote the formation of an active NRG1 resistosome. These possibilities are consistent with observations that *At*NRG1 associates with *At*EDS1 and *At*SAG101 to form a functional signalling complex in the immune-activated state (Lapin et al., 2019; Sun et al., 2021) and upon activation of PTI and TIR-NLR mediated ETI, a small proportion of *At*NRG1 forms stable resistosomes with *At*EDS1 and *At*SAG101(Feehan et al., 2023).

An unexpected observation here was that high expression of *Nb*NRG1 could inhibit TIR-NLR mediated cell death in *N. benthamiana* (**Supp Figure 6B**). Given that *Nb*NRG1 associates with *Nb*SAG101b in competition with *Nb*EDS1, we hypothesise that overexpression of *Nb*NRG1 may sequester *Nb*SAG101b away from the *Nb*EDS1/*Nb*SAG101b complex required to transduce the cell death signal from the TIR proteins to *Nb*NRG1. A recent discovery showed that the *Arabidopsis* NRG1C, a N-terminally truncated member of the NRG1 family, inhibits TIR-NLR induced cell death by antagonizing the function of full length NRG1A and NRG1B, possibly by interfering with the EDS1-SAG101-NRG1A/B axis (Wu et al., 2022). Interestingly, the NB, LRR and the combined NB-LRR domains of *Nb*NRG1 could interact with *Nb*SAG101b and functional chimeras containing its CEP domain (**Figure 7**). However, the isolated NB and LRR domains of *Nb*NRG1 showed a loss of specificity in that they could also interact with all the *Nb*EDS1 family and chimeric proteins. Thus, it seems that the NB and LRR domains act together to provide the specificity that allows NRG1 to receive signal only from *Nb*SAG101b.

### Model for signal transduction in TIR-EDS1/SAG101b/NRG1 pathway

**Supp Figure 17** shows a hypothetical model of TIR signalling events leading to activation of *Nb*NRG1. Effector recognition leads to a TIR-NLR forming a tetrameric resistosome which activates the TIR domain catalytic activity (Horsefield et al., 2019; Huang et al., 2022; Jia et al., 2022; Ma et al., 2020; Martin et al., 2020). The TIR domains associate with the NLP domains of EDS1 family heterodimers, to allow efficient transfer of signalling molecules, alter enzymatic activity and/or provide access to a substrate pre-bound to the EDS1 protein complex. After detection of the TIR-derived molecular signal, The EDS1/SAG101 complex undergoes a conformational shift allowing the CEP domain of SAG101 to interact with the NB and LRR domains of NRG1. This interaction results in activation of the *Nb*NRG1 protein likely through oligomerisation into a resistosome structure and formation of a Ca^2+^ permeable cation channel (Bi et al., 2021; Feehan et al., 2023; Jacob et al., 2021).

## Methods

### Generation of constructs

Details of primers and constructs used here are shown in Table S1-S3. All new constructs were generated by Gateway cloning (GWY; Invitrogen), with Phusion High-Fidelity DNA Polymerase (Thermo Fisher) used for PCR amplification of fragments flanked by attB sites for recombination into pDONR207 (BP reaction) and then into destination vectors (LR reaction). The binary vector pAM-PAT-35s-GWY-YFPv (or with 3xHA or 3xMyc tags) (Bernoux et al., 2008) was used for all *in planta* cell death and CoIP experiments, except for *Nb*NRG1 and *Nb*ADR1 for which the pBIN19-GWY-CFP (or 3xHA tag) vector (Cesari et al., 2016) was also used. The TIR-NLRC4 fusion constructs were previously described (Duxbury et al., 2020). For Split-Luciferase assays, genes were cloned into the binary vectors pDEST-GWY-nLUC and pDEST-GWY-cLUC (Gehl et al., 2011; Saur et al., 2019). For yeast two-hybrid and three-hybrid experiments, cDNAs were cloned into Gateway-compatible yeast two-hybrid vectors based on pGADT7 and pGBKT7 (Clontech) or into the pAG416GPD-ccdB-HA vector (Alberti et al., 2007) (pAG416GPD-ccdB-HA was a gift from Susan Lindquist (Addgene plasmid # 14244; http://n2t.net/addgene:14244; RRID: Addgene_14244)). Mutations were generated by DpnI-mediated site-directed mutagenesis (Stratagene). For plant two-hybrid assays, genes were cloned into the binary vectors pAM-PAT-GWY-BD and pAM-PAT-VP16-GWY (Chen et al., 2023).

### Transient expression in *N. benthamiana*

*N. benthamiana* plants carrying CRSIPR/Cas9-generated mutants were described previously [*eds1* (Qi et al., 2018), *eds1/pad4*, *pad4/sag101a/sag101b* (Gantner et al., 2019)] and *nrg1* was provided by J. Stuttmann. Plants were grown at 23°C with a 16-hours light period. For leaf infiltration, constructs were transformed into *Agrobacterium tumefaciens* strain GV3103_pMP90 (pAM-PAT constructs) or GV3101_pMP90 (pBIN19 and pDEST-GWY-nLUC/cLUC constructs) by electroporation. Cultures were grown at 28°C with shaking overnight in Luria-Bertani liquid medium containing 25 μg/ml of rifampicin, 15 μg/ml gentamicin and either 50 μg/ml of kanamycin (pBIN19 and pDEST vectors) or 25 μg/ml of carbenicillin (pAM-PAT vectors). Cells were harvested by centrifugation, resuspended in infiltration buffer (10 mM MES pH 5.6, 10 mM MgCl2 and 150 µM acetosyringone) and incubated at room temperature for 2 hours. The final bacterial concentration for cell death complementation and CoIP assays was OD600 = 0.5. Three to four individual plants were infiltrated for each combination of constructs and experiments were repeated at least three times. Plants were infiltrated 3-4 weeks after transplanting in the third and fourth leaves counting from the youngest apical leaf, which give the most robust cell death responses under our conditions. However, in *nrg1* mutant plants the 3^rd^/4^th^ leaves consistently showed a non-specific cell death response to *Agrobacterium* infiltration, so we used the 5^th^ and 6^th^ leaves which did not show non-specific cell death but gave a generally weaker response than the 3^rd^/4^th^ leaves of wild-type plants. Leaves were scanned 3-5 days after infiltration to record cell death responses.

### Yeast two-hybrid and yeast three-hybrid assays

Transformation of *Saccharomyces cerevisiae* strain HF7c was performed as described in the Yeast Protocols Handbook (Clontech), with transformants selected on synthetic dropout (SD) medium lacking tryptophan and leucine (-WL) to select for pGADT7 and pGBKT7 plasmids, or additionally lacking Uracil (-WLU) to select for pAG416GPD vectors. To assess protein interactions, three independent colonies were grown on medium lacking leucine, tryptophan, and histidine (-HWL two-hybrid) or also lacking uracil (-HWLU, three-hybrid) at 30°C for 3-4 days.

### Split-luciferase and plant two-hybrid assays

The split-luciferase (Split-LUC) complementation assay was performed as previously described (Saur et al., 2019). Three disks (0.38 cm diameter) from three independent leaves were harvested into 1.5 ml tube two days after infiltration at 4-week-old *N. benthamiana* plants. Samples were ground to fine powder using pestles on dry ice and then 100 μl 2x cell culture lysis buffer was added (Promega, E1531; with 150 mM Tris-Hcl pH7.5). 50 μl of leaf extract was mixed with 50 μl luciferase substrate (Promega, E1501) in a white-bottomed 96-well plate, and the light emission was measured using a microplate luminometer (Fluostar Omega) at 1 s/well. The final Agrobacterium concentration of each construct was OD600=1.5, except for the pDEST-NLP-*Nb*EDS1-cLUC, pDEST-*Nb*SAG101b-cLUC and pDEST-*Nb*PAD4-cLUC, which was OD600=0.15 as these constructs showed higher protein expression than other pDEST constructs. For plant two-hybrid assays, Ruby expression was detected as described previously (Chen et al., 2023). Four-week-old *N. benthamiana* plants were used for the infiltration of Agrobacteria containing UAS-Ruby and TIR/EDS1 constructs. Whole leaves were harvested 3-5 days after infiltration and scanned and then cleared using ethanol. Three disks from three independent decolourised leaves were collected in a 1.5 ml tube with 0.5 ml water to extract betalain. The absorbance of betalain was measured at 538nm. The Agrobacterium concentration was OD600=0.2 for the pAM-TIR-BD constructs, and OD600=0.8 for the pAM-VP16-*Nb*EDS1 constructs.

### Protein Extraction, Immunoblot, and Coimmunoprecipitation

For immunoblot analysis, proteins were extracted from plant tissues directly into loading buffer (0.125 M Tris-HCl, pH 7.5, 4% SDS, 20% Glycerol, 0.2 M DTT, 0.02% Bromophenol Blue) while yeast protein extraction was performed following a post-alkaline method (Kushnirov, 2000). For Co-immunoprecipitation experiments agrobacterium cultures were infiltrated at OD600 = 0.5 and proteins were extracted as previously described (Cesari et al., 2013). For immunodetection, proteins were separated by SDS-PAGE and transferred to nitrocellulose membranes which were blocked in. 5% skimmed milk. Membranes were incubated with Anti-HA-Peroxidase, High Affinity (Roche, REF 12013819001) or mouse anti-GFP (Roche, REF 11814460001) and anti-Myc antibodies (Roche, REF 11667149001) followed by goat anti-mouse antibodies conjugated with horseradish peroxidase (Biorad, REF 170–5047), or polyclonal rabbit anti-luciferase (Sigma-Aldrich, REF L0159), anti-GAL4BD (Sigma-Aldrich, REF G3042) and anti-VP16 (Sigma-Aldrich, REF V4388) antibodies followed by anti-rabbit horseradish peroxidase (Sigma) to detect -HA, YFP, Myc, nLUC, cLUC, GAL4BD and VP16 tagged proteins respectively. Signals were detected using the SuperSignal West Femto chemiluminescence kit (Pierce). Ponceau S was used to stain membranes to confirm equal protein loading.

---

**Table S1.**
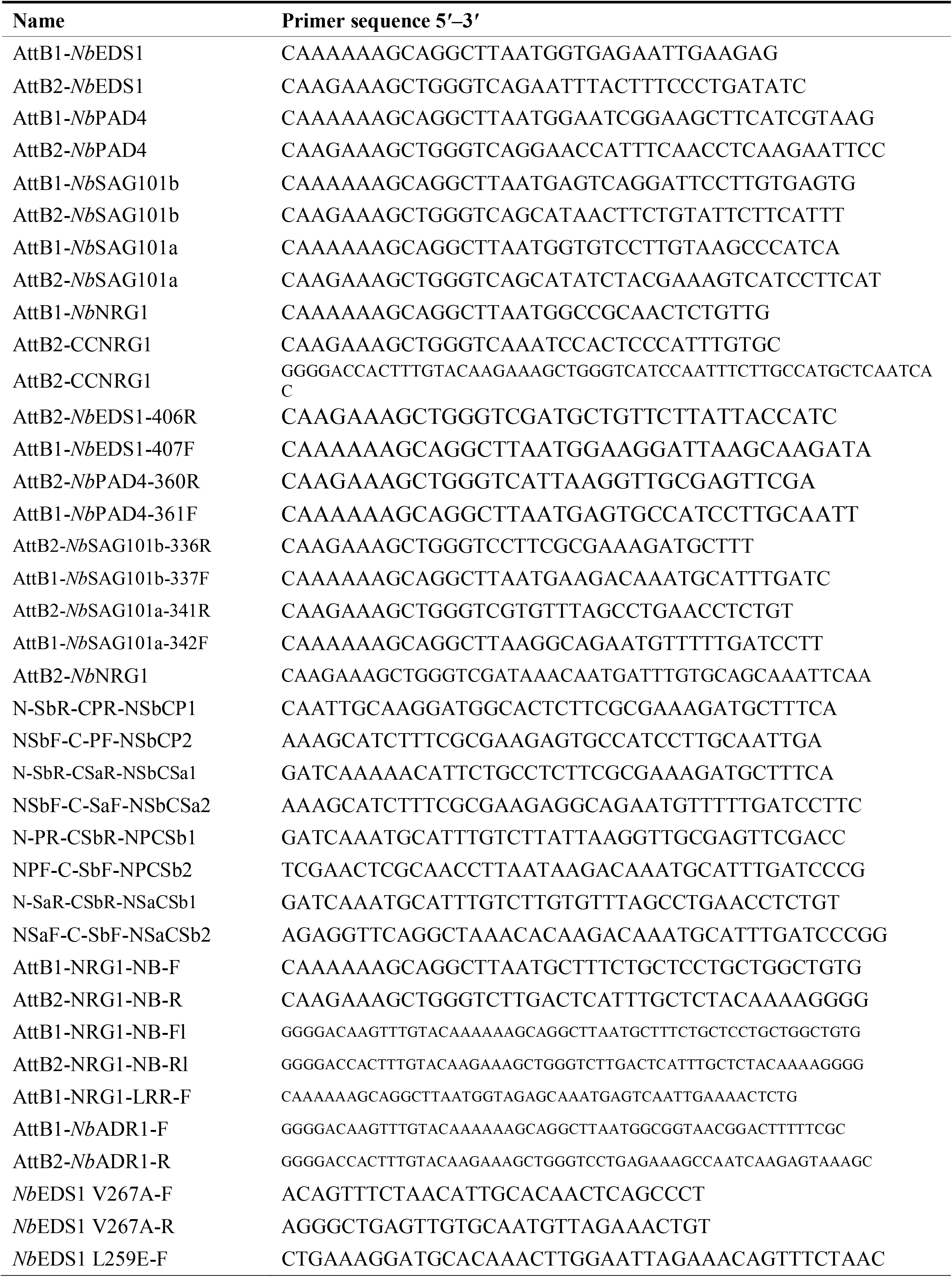
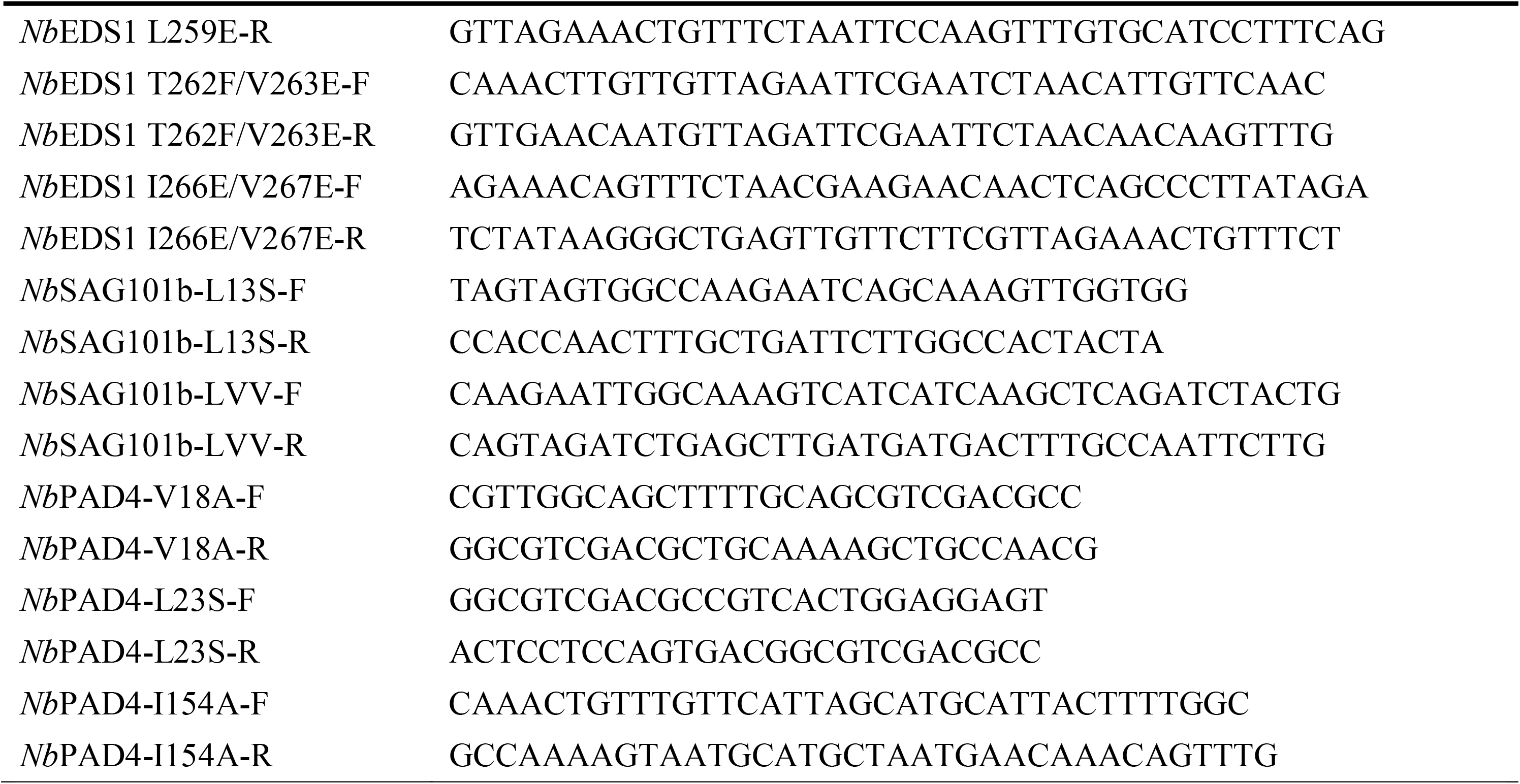
Primers used in this study.

**Table S2.** Constructs used in this study for yeast assays.

| Use | DNA insert | Plasmid backbone | Cloning method/Source |
| --- | --- | --- | --- |
| Constructs for yeast assays | *NbEDS1* | pGADT7-GWY | Gateway |
|  | *NbEDS1* | pGBK7-GWY | Gateway |
|  | *NbEDS1* | pAG416GPD-GWY | Gateway |
|  | *NbPAD4* | pGADT7-GWY | Gateway |
|  | *NbPAD4* | pGBK7-GWY | Gateway |
|  | *NbPAD4* | pAG416GPD-GWY | Gateway |
|  | *NbSAG101a* | pGADT7-GWY | Gateway |
|  | *NbSAG101a* | pGBK7-GWY | Gateway |
|  | *NbSAG101b* | pGADT7-GWY | Gateway |
|  | *NbSAG101b* | pGBK7-GWY | Gateway |
|  | *NbSAG101b* | pAG416GPD-GWY | Gateway |
|  | d28L6TIR (29-233) | pGADT7-GWY | (Bernoux et al., 2011) |
|  | d28L6TIR (29-233) | pGBK7-GWY |  |
|  | RPS4TIR (1-183) | pGADT7-GWY | (Zhang et al., 2017a) |
|  | RPS4TIR (1-183) | pGBK7-GWY |  |
|  | SNC1TIR (1-226) | pGADT7-GWY | Gateway |
|  | SNC1TIR (1-226) | pGBK7-GWY | Gateway |
|  | Run1TIR (1-198) | pGADT7-GWY | Gateway |
|  | Run1TIR (1-198) | pGBK7-GWY | Gateway |
|  | NLP-NbEDS1 (1-406) | pGBK7-GWY | Gateway |
|  | CEP-NbEDS1 (407-607) | pGBK7-GWY | Gateway |
|  | NLP-NbPAD4 (1-360) | pGADT7-GWY | Gateway |
|  | CEP-NbPAD4 (361-609) | pGADT7-GWY | Gateway |
|  | NLP-NbSAG101b (1-336) | pGADT7-GWY | Gateway |
|  | CEP-NbSAG101b (337-581) | pGADT7-GWY | Gateway |
|  | *NbEDS1* LTVIV | pGADT7-GWY | Gateway |
|  | *NbEDS1* LTVIV | pGBK7-GWY | Gateway |
|  | *NbEDS1* LTVIV | pAG416GPD-GWY | Gateway |
|  | *NbPAD4* VLI | pGADT7-GWY | Gateway |
|  | *NbPAD4* VLI | pGBK7-GWY | Gateway |
|  | *NbPAD4* VLI | pAG416GPD-GWY | Gateway |
|  | *NbSAG101b* LLVV | pGADT7-GWY | Gateway |
|  | *NbSAG101b* LLVV | pGBK7-GWY | Gateway |
|  | *NbSAG101b* LLVV | pAG416GPD-GWY | Gateway |

**Table S3.**
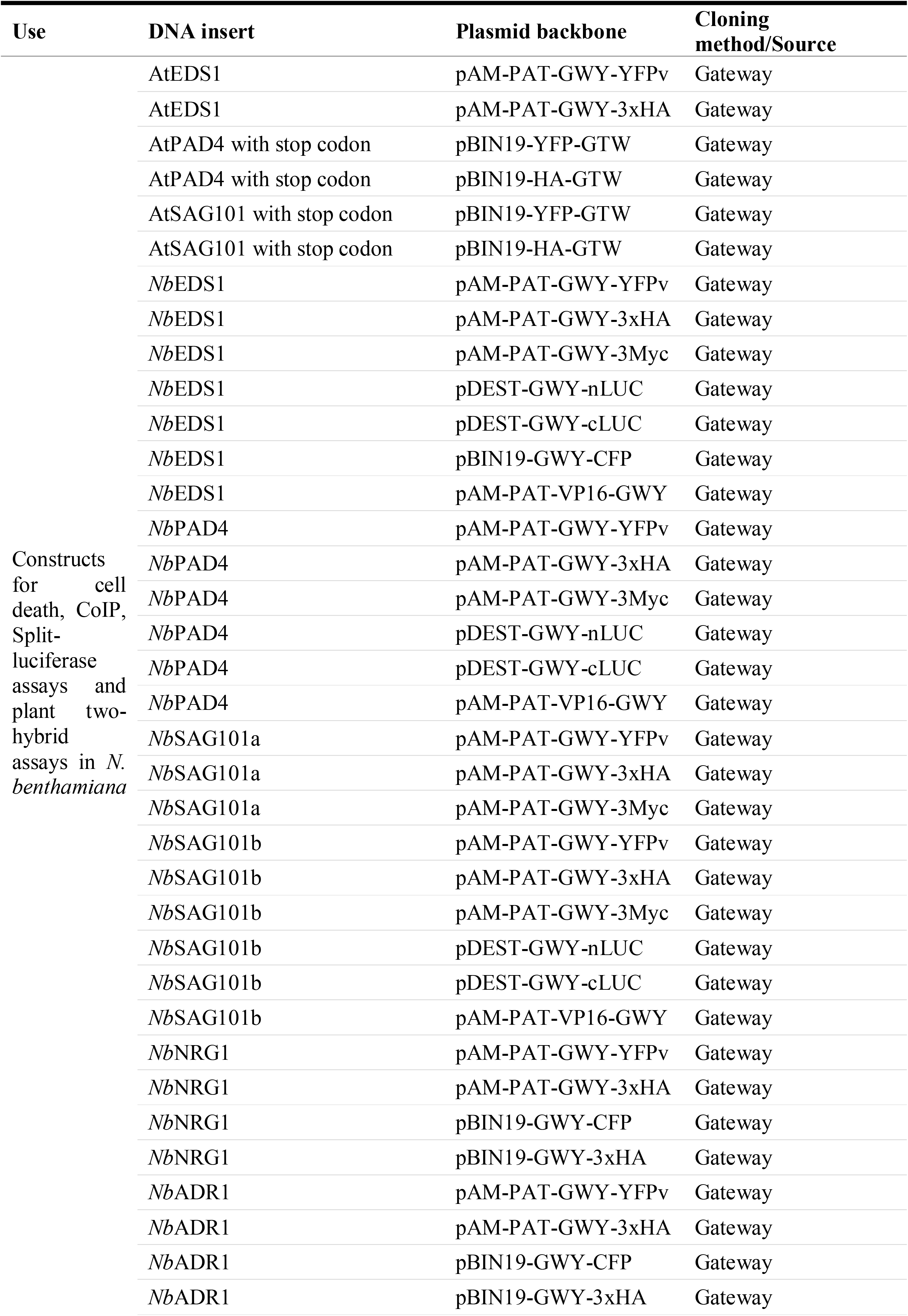
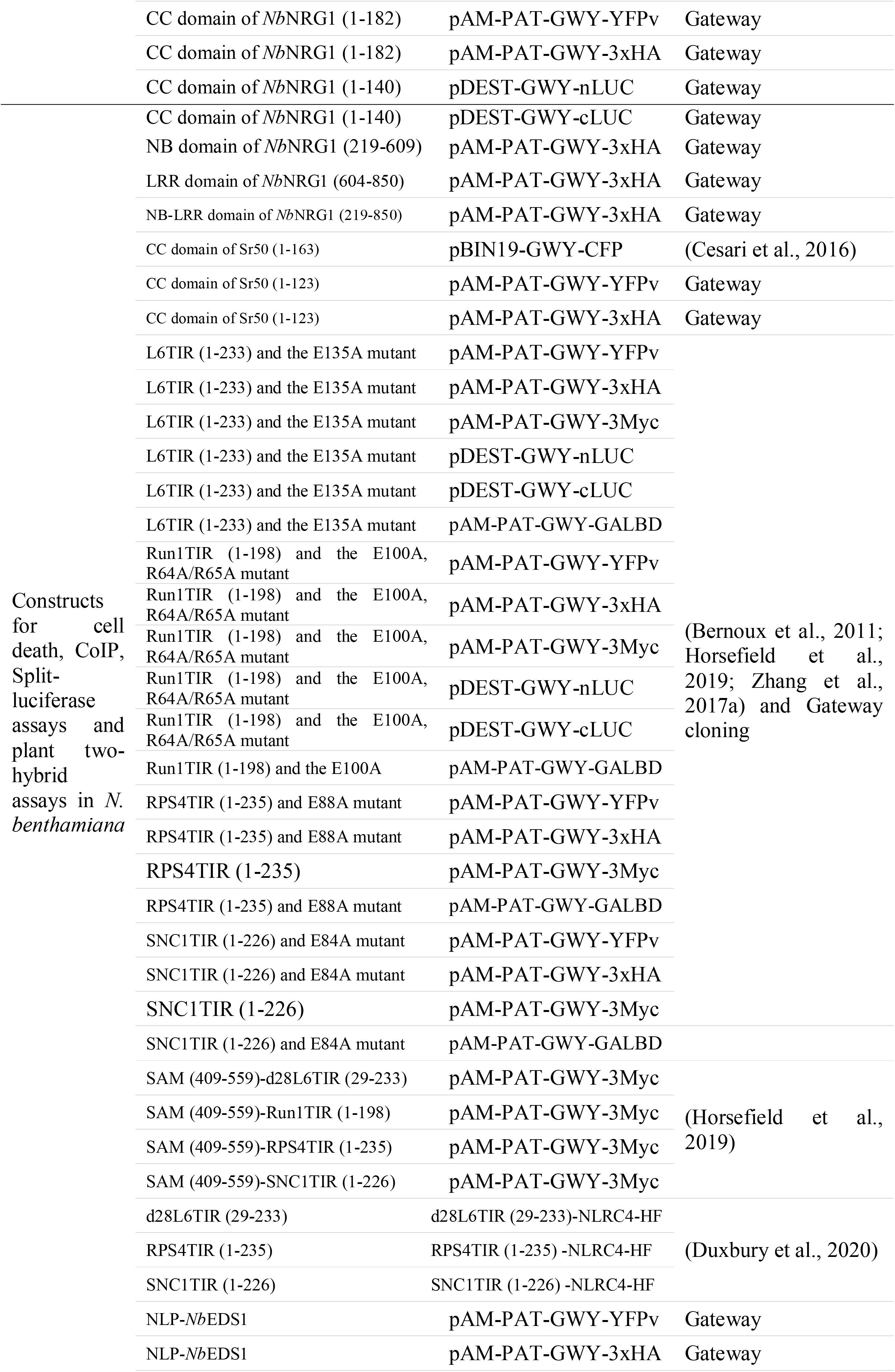
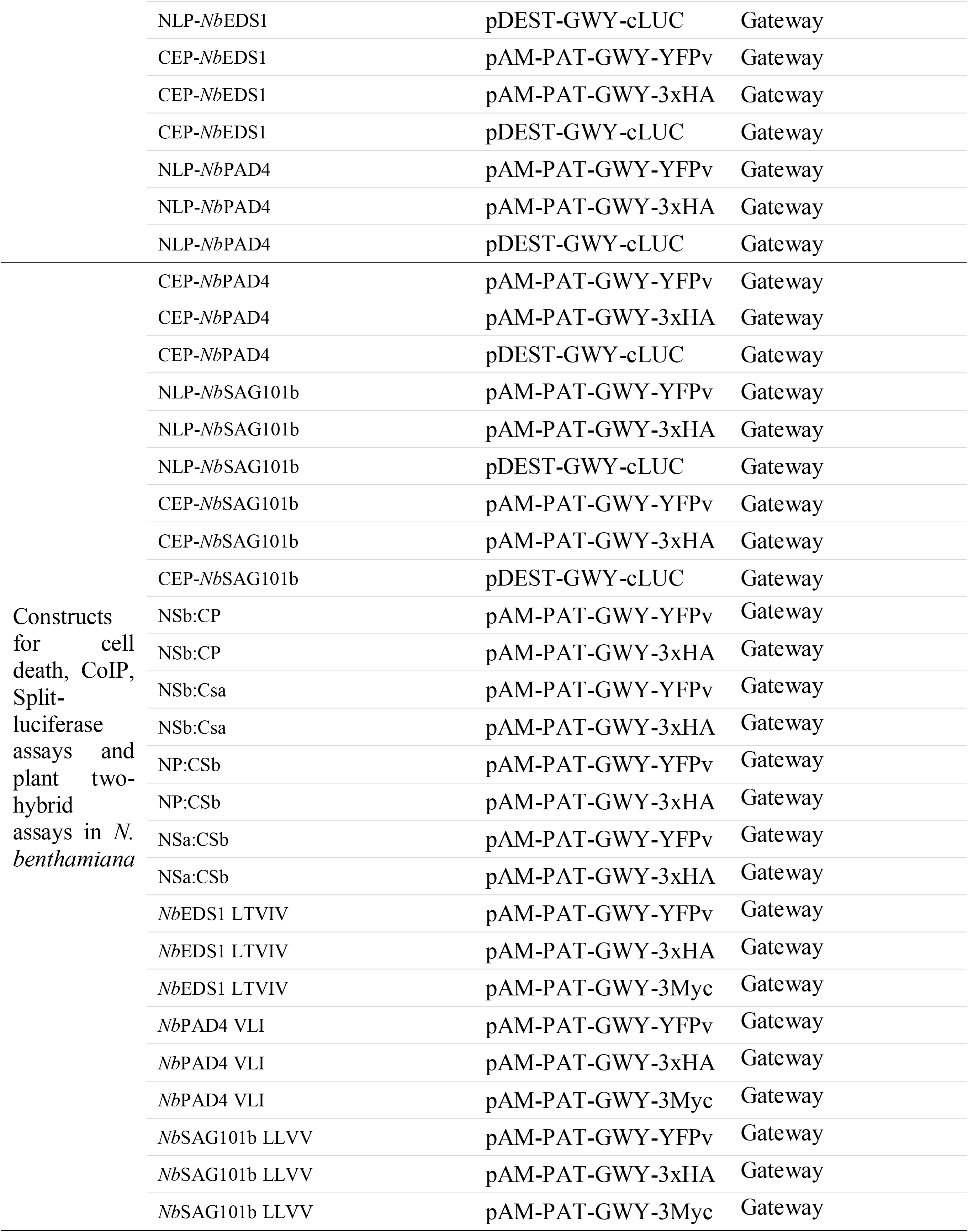
Constructs used in this study for *in planta* assays.

## Author contributions

JC conducted experimental work and drafted the manuscript. PND, JPR, MB conceived the study. All authors contributed to data analysis and interpretation and manuscript writing.

## Acknowledgments

The mutant lines of N. benthamiana were kindly provided by B. Staskawicz (*eds1*) and J. Stuttmann (*eds1/pad4*, *pad4/sag101a/sag101b* and *nrg1*)

## Supplemental Figure Legends

**Supplemental Figure S1. TIR alone, TIR fusion and full length TIR-NLR constructs used for cell death assays.** L6TIR, Run1TIR, SNC1TIR and RPS4TIR fused with YFP tag are autoactive in wildtype *N. benthamiana*. Full length flax resistance proteins L6 and M induce strong HR when transiently expressed together with effector proteins AvrL567 and AvrM, respectively, possibly by formation of a tetrameric resistosome as observed for ROQ1 and RPP1. L6 MHV, M MHV and M1 MHV are full length TIR-NLRs contain D to V mutations in the conserved MHD motif that result in strong autoactive cell death induction *in planta*. d28L6TIR is a shorter non-autoactive version of L6TIR without the N-terminal 28 amino acid Golgi membrane anchoring sequence (Bernoux et al., 2023). Run1TIR, SNC1TIR, RPS4TIR and d28L6TIR fused to the oligomerising SAM domain of SARM1 are strongly autoactivity (Horsefield et al., 2019), while fusions to NLRC4 can be activated to induce cell death by co-expression with NAIP5 and FlaA (Duxbury et al., 2020). pTIR: plant TIRs; NB: nucleotide-binding; LRR: leucine-rich repeat; CARD: caspase activation and recruitment domain; NACHT: NACHT, NAIP, CIITA, HET-E, TP1; SAM: sterile alpha motif.

**Supplemental Figure S2. *Nb*EDS1 is required for TIR-mediated cell death in *N. benthamiana*.** Complementation of TIR-mediated cell death in the *eds1* mutant line. Indicated proteins were expressed in wild type *N. benthamiana* (*N.b*) or in *eds1* mutant lines alone or in combination with *Nb*EDS1 fused with a 3xHA tag by Agrobacterium-mediated transient expression. Photos were taken at 5 dpi.

**Supplemental Figure S3. RPS4TIR constructs have residual cell death activity in *eds1*.** RPS4TIR fused with a YFP tag or to the SAM domain with 3xMyc tag induced strong cell death in wildtype *N.b*, but only a weak residual cell death in *eds1*. Introduction of the NADase catalytic site mutation E88A (RPS4TIR-YFP E88A and SAMRPS4TIR-3Myc E88A) or the non-oligomerising SAM5M fusion (SAM5MRPS4TIR-3Myc) abolished it. The CC domain of Sr50 (CCSr50-YFP) was used as the positive control. Photos were taken at 5 dpi.

**Supplemental Figure S4. *Nb*PAD4 is not required for TIR-mediated cell death in *N. benthamiana*.** Complementation of TIR-mediated cell death in *eds1pad4* mutant line. Indicated proteins were expressed in wild type *N. benthamiana* (*N.b*) or in the *eds1pad4* double mutant line either alone or in combination with *Nb*EDS1 or *Nb*PAD4 or both by Agrobacterium-mediated transient expression. *Nb*EDS1 and *Nb*PAD4 were fused with 3xHA tag. Photos were taken at 5 dpi.

**Supplemental Figure S5. *Nb*SAG101b is required for TIR-mediated cell death in *N. benthamiana*. *Nb*PAD4 and *Nb*SAG101a are not necessary.** Complementation of TIR-mediated cell death in the *pss* triple mutant line after Agrobacterium-mediated transient expression. Indicated proteins were expressed in wild type *N. benthamiana* (*N.b*) or in the *pss* triple mutant line either alone or together with *Nb*PAD4, *Nb*SAG101a, *Nb*SAG101b fused with 3xHA tag or combinations of these. Photos were taken at 5 dpi.

**Supplemental Figure S6. *Nb*NRG1 is required for TIR-mediated cell death in *N. benthamiana*** (A). Cell death recovery in *nrg1* triple mutant line by Agrobacterium-mediated transient expression of *NbNRG1*. Indicated proteins that fused with YFP tag were expressed alone or together with *Nb*NRG1 that fused with 3xHA tag. Photos were taken at 5 dpi. (B). *Nb*NRG1 inhibited autoactive TIRs induced cell death in WT plants (Infiltration of *Nb*NRG1; OD600 ≥ 0.5). Photos were taken at 5 dpi. Western blot shows the accumulation of TIR proteins with and without *Nb*NRG1.

**Supplemental Figure S7. Overexpression of *Nb*NRG1 and *Nb*ADR1 fused to 3xHA tag induce cell death in *N. benthamiana.*** Autoactivity of *Nb*NRG1 and *Nb*ADR1 expressed from pAM-PAT-35s-GWY-3xHA or pBIN19-35s-GWY-3xHA vectors tested in WT and mutant *N. benthamiana* plants (OD600 = 2.0). Western blot showed the accumulation of *Nb*NRG1, *Nb*ADR1 and *Nb*EDS1 proteins from each vector.

**Supplemental Figure S8. CC-NLRs and autoactive CC domains induce cell death in *N. benthamiana* mutant lines.** Cell death was tested in *N. benthamiana* WT, *eds1*, *eds1pad4*, *pss* and *nrg1* mutant lines. Full length Sr50 (OD600 = 0.1) was co-infiltrated with YFP-AvrSr50 (OD600 = 0.5). CCSr50-YFP (OD600 = 0.5) is the autoactive CC domain from Sr50. CCNRG1-YFP is the autoactive CC domain from *Nb*NRG1. pAM*-Nb*NRG1-3xHA (OD600= 0.5) is not autoactive due to low expression from the construct.

**Supplemental Figure S9. Plant TIRs interact with *Nb*EDS1.** (A). CoIP showed L6TIR-3xHA interacts with *Nb*EDS1-YFP, *Nb*PAD4-YFP and *Nb*SAG101b-YFP. TIRs-3xHA in combination with YFP alone and TIRs-YFP in combination with CC domain of Sr50 that fused to 3xHA tag were used as negative controls. (B). CoIP of *Nb*EDS1-3xHA with wildtype Run1TIR-YFP, Run1TIR-YFP E100A (NADase catalytic site mutations) and Run1TIR-YFP RRAA (R64AR65A), the combinations were transiently expressed in *N. benthamiana* leaves. (C). CoIP of *Nb*EDS1-3xHA with wildtype L6TIR-YFP and AE/DE interfaces mutations, the combinations were expressed in *pss* mutant line. (D). CoIP of *Nb*EDS1-3xHA with plant TIRs-YFP. The combinations were transiently expressed in both wildtype and *pad4/sag101a/sag101b* mutant plants. (E). Competition CoIP experiments of *Nb*EDS1-3xHA with TIR-YFP proteins with and without *Nb*SAG101b-3Myc.

**Supplemental Figure S10. Plant TIRs interact with *Arabidopsis* EDS1, PAD4 and SAG101 in CoIP.** (A) CoIP experiment of *At*EDS1 and different TIRs. The combinations of *At*EDS1-3xHA/-YFP with wildtype TIR-YFP/-3xHA were transiently expressed in *pss* mutant plants. (B) CoIP experiment of *At*PAD4*, At*SAG101 with different plant TIRs in *eds1* plants. *At*PAD4*, At*SAG101 were fused to N-terminal 3xHA tag, TIRs were fused to a YFP tag in the C-terminal of TIR proteins. Leaf samples were collected 1 dpi.

**Supplemental Figure S11. Functional assays of split-luciferase constructs in *N. benthamiana*.** (A). Cell death of L6TIR and Run1TIR fused with cLUC and nLUC in *N. benthamiana*. (B). Cell death complementation assays of L6TIR and Run1TIR fused with YFP tags by *Nb*SAG101b and *Nb*EDS1 fused with cLUC and nLUC in *pss* and *eds1* mutant lines. (C). Protein accumulation from expression of -nLUC and -cLUC fusion constructs. All constructs were transiently expressed in *N. benthamiana* plants. Leaf samples were collected 2 dpi. Total proteins were extracted and separated by SDS-PAGE. Western blot membranes were incubated with an anti-luciferase antibody. -cLUC indicates fusion to the C-terminal fragment of luciferase, -nLUC indicates fusion to the N-terminal fragment of luciferase. (D) Split-LUC assays of NbEDS1, NbSAG101b, NbPAD4 sub-domains fused to cLUC with Run1TIR-nLUC. cLUC indicates C-terminal fragment of luciferase, nLUC indicates the N-terminal fragment of luciferase, NLP indicates N-terminal lipase-like domain, CEP indicates C-terminal EP domain. Leaf samples were taken at 2 dpi. The significant difference of luciferase with Run1TIR-nLUC between NLP and CEP domains were labelled with asterisk (***: p<0.001). The combinations of TIR-nLUC+EDS1-cLUC were infiltrated in the *pss* mutant plants, the other combinations were infiltrated in the *eds1* mutant plants.

**Supplemental Figure S12. Protein accumulation for expression of plant two-hybrid constructs.** All constructs were transiently expressed in *N. benthamiana* plants, the TIR constructs were infiltrated in *eds1* mutant plants. Leaf samples were collected 2 dpi. Total proteins were extracted and separated by SDS-PAGE. Western blot membranes were incubated with an anti-GAL4BD or anti-VP16 antibodies.

**Supplemental Figure S13. Tests for interaction between plant TIRs and *Nb*EDS1 family proteins by yeast two-hybrid.** (A). Yeast two-hybrid interaction assays between *Nb*EDS1 family proteins fused to the AD domain and plant TIRs fused to the BD domain. (B). Yeast two-hybrid interaction assay between *Nb*EDS1 family proteins fused to the BD domain and plant TIRs fused to the AD domain. *Nb*EDS1-BD+ *Nb*PAD4-AD was used as positive control.C). Heterodimers of full length and truncated *Nb*EDS1 family proteins in Y2H. The full length and truncated *Nb*EDS1 proteins were fused to the BD domain, the full length and truncated *Nb*SAG101b and *Nb*PAD4 proteins were fused to the AD domain. NLP indicates N-terminal lipase-like domain, CEP indicates C-terminal EP domain. Yeast colonies were diluted and grown on media lacking tryptophan and leucine (-WL, growth control), or additionally lacking histidine (-WLH, interaction selection). Pictures were taken after 3 days of growth at 30LJ.

**Supplemental Figure S14. *Nb*EDS1-BD activated yeast HIS3 reporter gene expression when expressed with free *Nb*SAG101b and *Nb*PAD4.** A). Yeast two-hybrid interaction assay for *Nb*EDS1-BD with free *Nb*SAG101b and *Nb*PAD4 variants and *Nb*SAG101b-BD with free *Nb*EDS1. (B). Yeast three-hybrid assay of AD domain in combination with *Nb*EDS1-BD and *Nb*SAG101b-BD with free *Nb*EDS1, *Nb*SAG101b and *Nb*PAD4 variants. Yeast colonies were diluted and grown on media lacking tryptophan and uracil (-WU, growth control) and media lacking tryptophan, leucine and uracil (-WLU, growth control), or additionally lacking histidine (-HWU, growth control) and (-HWLU, interaction selection). Pictures were taken after 3 days of growth at 30LJ.

**Supplemental Figure S15. Constructs used in yeast assays have protein expression.** The expression of constructs used in yeast assays were detected by immunoblot analysis. AD fusion and ‘free’ proteins contain HA tags and were detected using Anti-HA antibody. BD fusion proteins contain a Myc tag and were detected using Anti-Myc antibody.

**Supplemental Figure S16. Homology modelling structures of *Nb*EDS1 heterocomplexes with chimeric proteins.** Modelling structures of *Nb*EDS1 heterocomplexes and domain swap chimeras. NSa: N-terminal lipase domain of *Nb*SAG101a; NP: N-terminal lipase domain of *Nb*PAD4; CSb: C-terminal domain of *Nb*SAG101b; CSa: C-terminal domain of *Nb*SAG101a; CP: C-terminal domain of *Nb*PAD4. Structures were modelled on the Arabidopsis *At*EDS1-*At*SAG101 crystal structure (PDB 4nfu) using SWISS-MODEL. Pictures were made using PyMOL (NSb: N-terminal lipase domain of NbSAG101b (aa 1-336); NSa: N-terminal lipase domain of NbSAG101a (aa 1-341); NP: N-terminal lipase domain of NbPAD4 (aa 1-360); CSb: C-terminal domain of NbSAG101b (aa 337-581); CSa: C-terminal domain of NbSAG101a (aa 342-583); CP: C-terminal domain of NbPAD4 (aa 361-609).

**Supplemental Figure S17. Model for signal transduction in TIR-EDS1/SAG101b/NRG1 pathway.** Proposed model for signal transduction in TIR-*Nb*EDS1/*Nb*SAG101/*Nb*NRG1 pathway. Monomeric TIR domains in inactive TIR-NLRs are capable of interacting with all EDS1 family proteins via their N-terminal lipase-like domains. Upon activation of the TIR-NLR, the active TIR oligomers assemble on the *Nb*EDS1 heterodimers, ensuring that the TIR catalytic function is active in close proximity to the EDS1/SAG101 heterodimer. Once the unstable small signalling molecules are generated they are bound the CEP pocket of the EDS1/SAG101 complex, inducing an allosteric change in the *Nb*SAG101b CEP domain. This allows interaction of the exposed CEP of *Nb*SAG101b with *Nb*NRG1 to activate *Nb*NRG1 leading to formation of the active *Nb*NRG1 resistosome.

