## supplementrary figures for "Plant TIR domains physically interact with EDS1 family proteins to propagate immune signalling"

**Supplemental Figure S1. TIR alone, TIR fusion and full length TIR-NLR constructs used for cell death assays.** L6TIR, Run1TIR, SNC1TIR and RPS4TIR fused with YFP tag are au

### Supplemental Figure 1

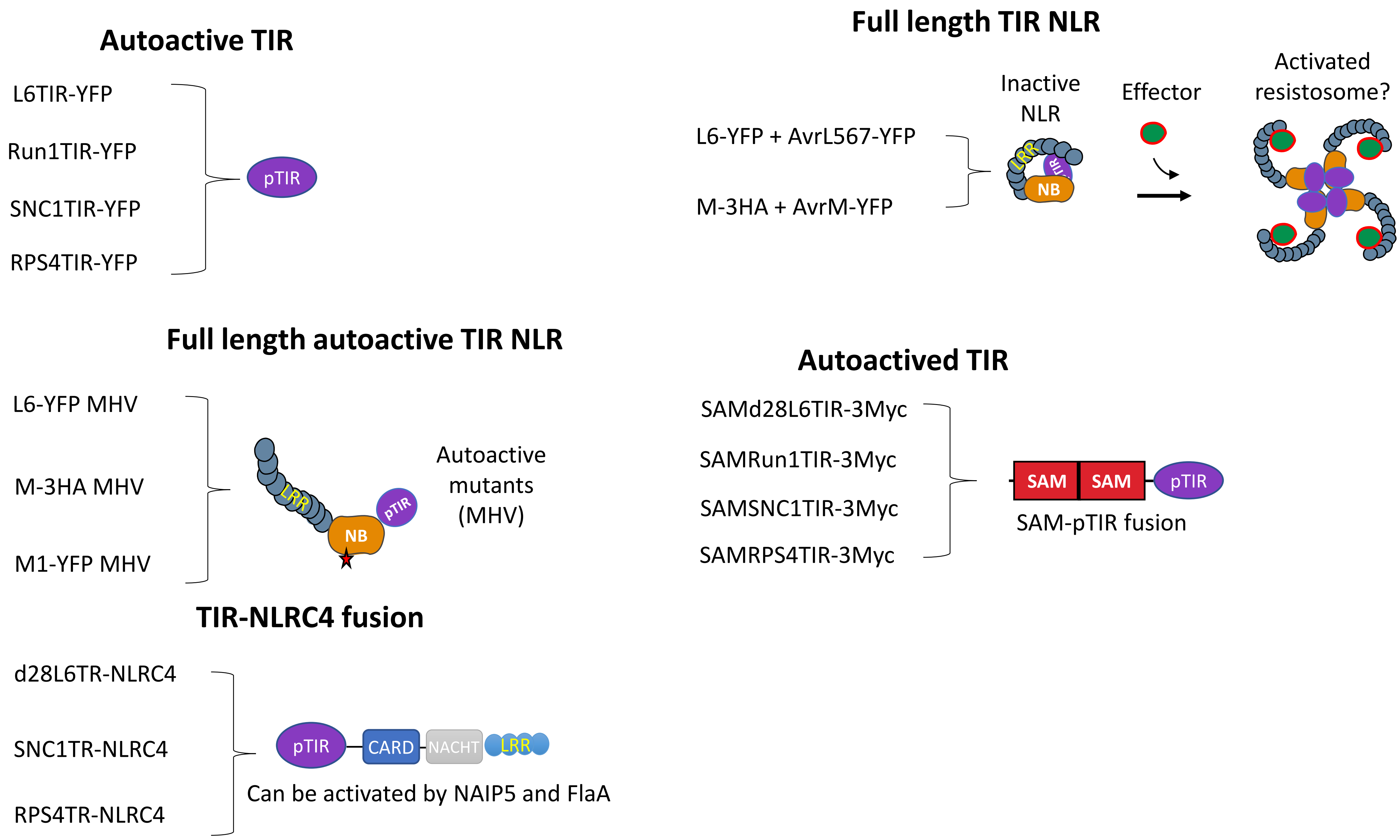


**Supplemental Figure S1. TIR alone, TIR fusion and full length TNL constructs used for cell death assay.**

L6TIR, Run1TIR, SNC1TIR and RPS4TIR fused with YFP tag are autoactive in wildtype *N. benthamiana*. Full length flax resistance protein L6 and M could induce strong HR when transient expression together with effector protein AvrL567 and AvrM. L6 MHV, M MHV and M1 MHV are autoactive full length TNLs, they can cause strong cell death *in planta*. d28L6TIR is a shorter non-autoactive version of L6TIR without the N-terminal 28 amino acid Golgi apparatus anchoring peptide. Oligomerisation of d28L6TIR by fusion with SAM or NLRC4, make the non-autoactive version of TIR autoactive. Run1TIR, SNC1TIR and RPS4TIR fused with either SAM or NLRC4 can cause strong cell death. pTIR: plant TIRs; NB: nucleotide-binding; LRR: leucine-rich repeat; CARD: caspase activation and recruitment domain; NACHT: NACHT, NAIP, CIITA, HET-E, TP1; SAM: sterile alpha motif.

#### Supplemental Figure 2

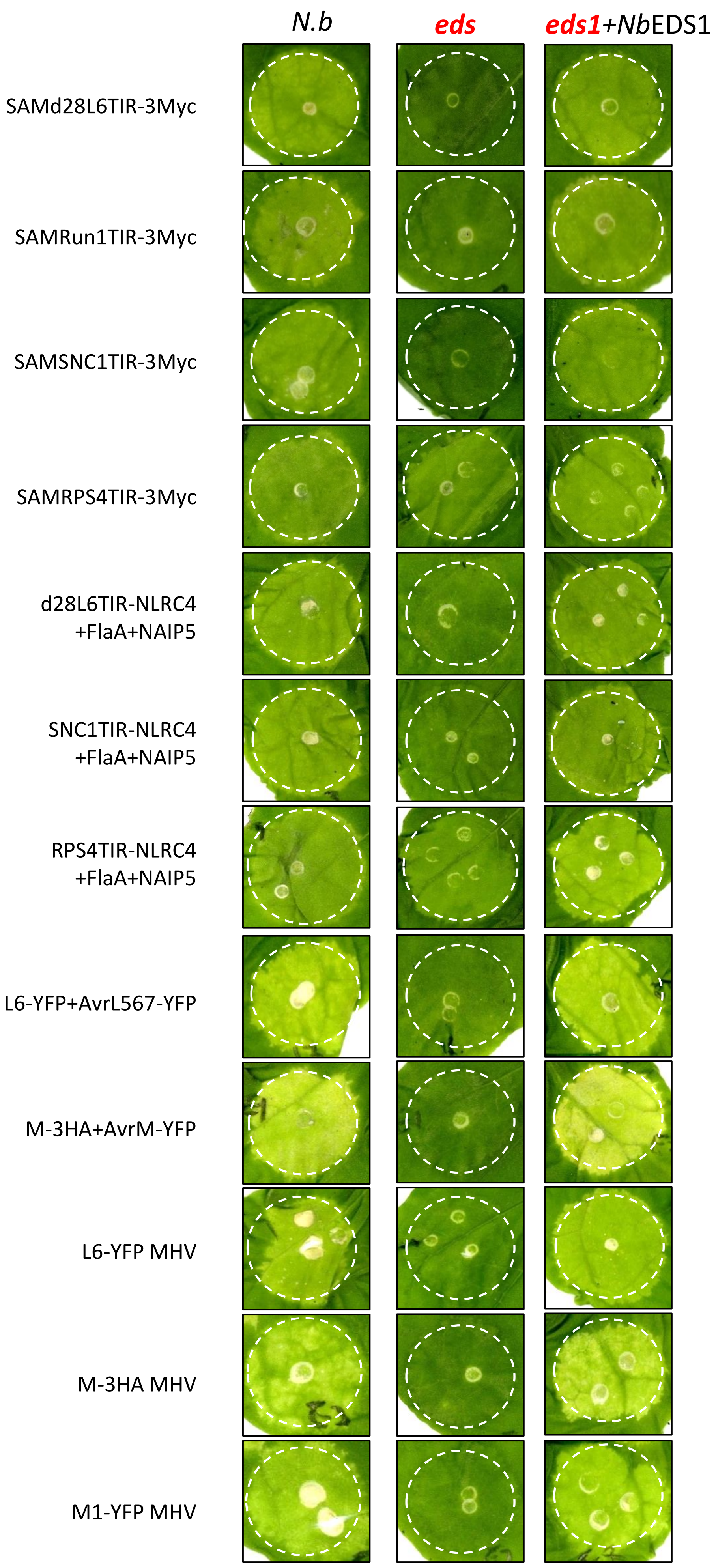


**Supplemental Figure S2. *Nb*EDS1 is required for TIR-mediated cell death in *N. benthamiana*.** Complementation of TIR-mediated cell death in the *eds1* mutant line. Indicated proteins were expressed in wild type *N. benthamiana* (*N.b*) or in *eds1* mutant lines alone or in combination with *Nb*EDS1 fused with a 3xHA tag by Agrobacterium-mediated transient expression. Photos were taken at 5 dpi.

### Supplemental Figure 3

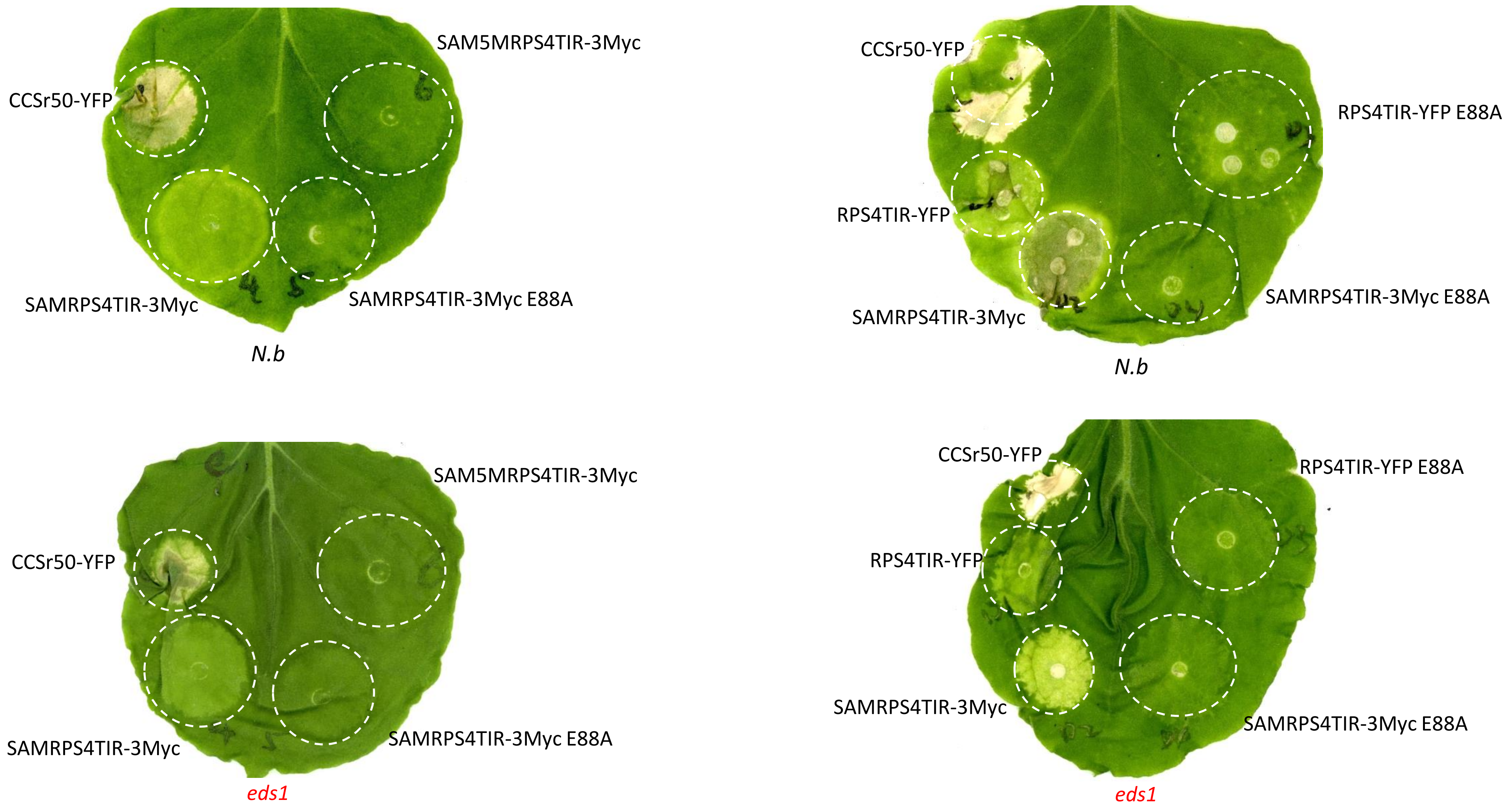


**Supplemental Figure S3. RPS4TIR constructs have residual cell death activity in *eds1*.** RPS4TIR fused with a YFP tag or to the SAM domain with 3xMyc tag induced strong cell death in wildtype *N.b*, but only a weak residual cell death in *eds1*. Introduction of the NADase catalytic site mutation E88A (RPS4TIR-YFP E88A and SAMRPS4TIR-3Myc E88A) or the non-oligomerising SAM5M fusion (SAM5MRPS4TIR-3Myc) abolished it. The CC domain of Sr50 (CCSr50-YFP) was used as the positive control. Photos were taken at 5 dpi.

### Supplemental Figure 4

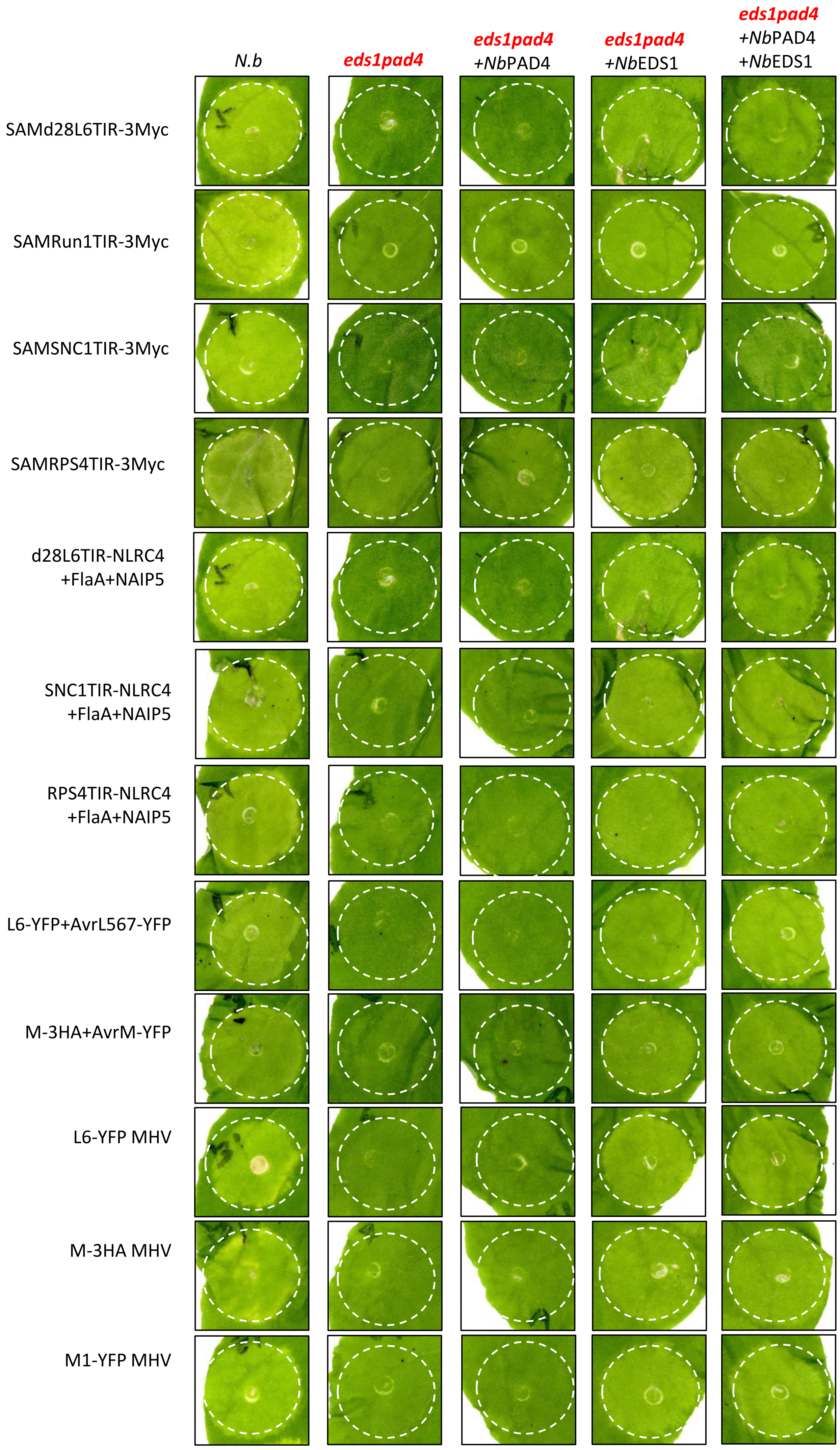


**Supplemental Figure S4. *Nb*PAD4 is not required for TIR-mediated cell death in *N. benthamiana*.** Complementation of TIR-mediated cell death in *eds1pad4* mutant line. Indicated proteins were expressed in wild type *N. benthamiana* (*N.b*) or in the *eds1pad4* double mutant line either alone or in combination with *Nb*EDS1 or *Nb*PAD4 or both by Agrobacterium-mediated transient expression. *Nb*EDS1 and *Nb*PAD4 were fused with 3xHA tag. Photos were taken at 5 dpi.

### Supplemental Figure 5

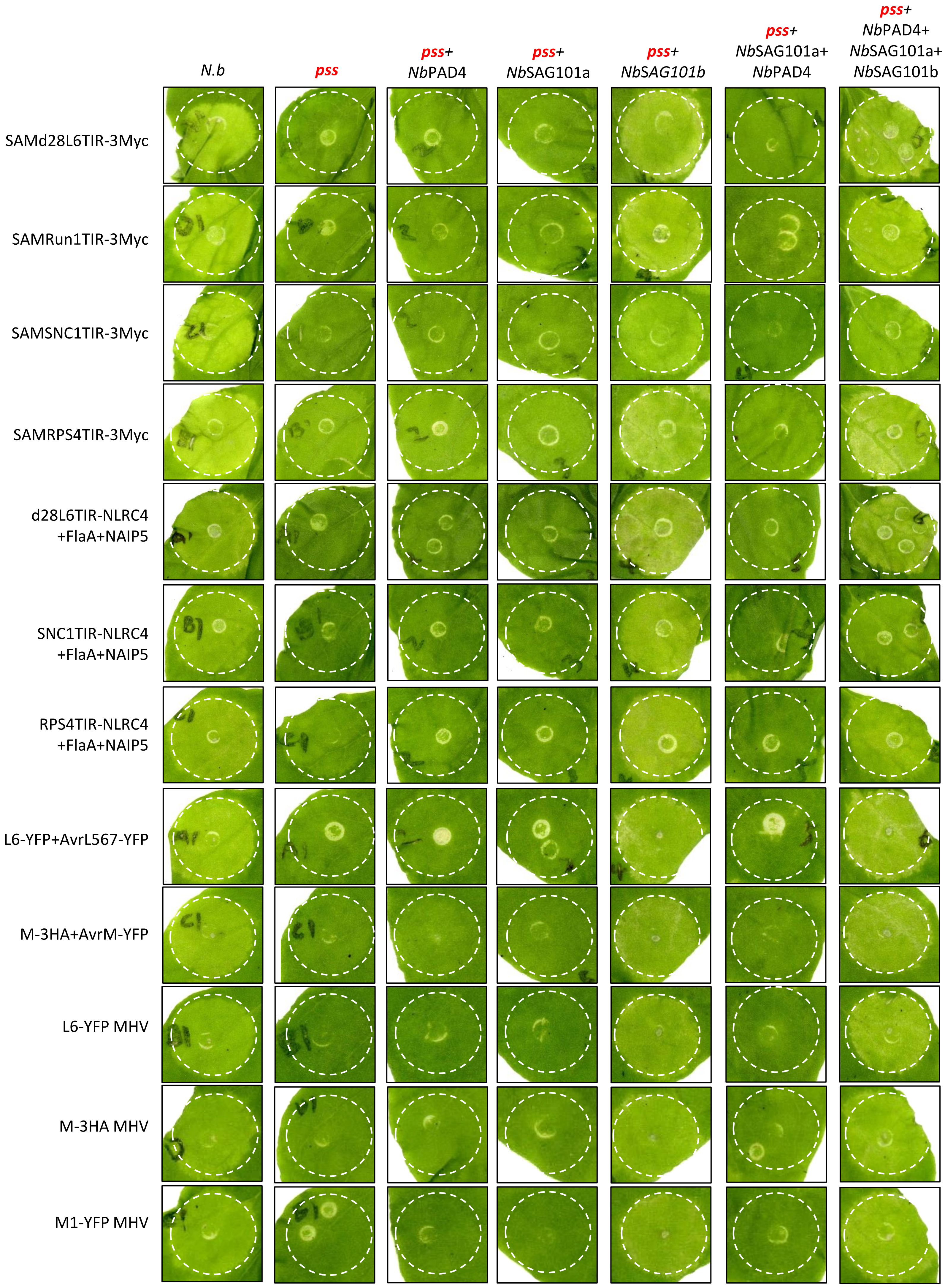


**Supplemental Figure S5. *Nb*SAG101b is required for TIR-mediated cell death in *N. benthamiana*. *Nb*PAD4 and *Nb*SAG101a are not necessary.** Complementation of TIR-mediated cell death in the *pss* triple mutant line after Agrobacterium-mediated transient expression. Indicated proteins were expressed in wild type *N. benthamiana* (*N.b*) or in the *pss* triple mutant line either alone or together with *Nb*PAD4, *Nb*SAG101a, *Nb*SAG101b fused with 3xHA tag or combinations of these. Photos were taken at 5 dpi.

### Supplemental Figure 6

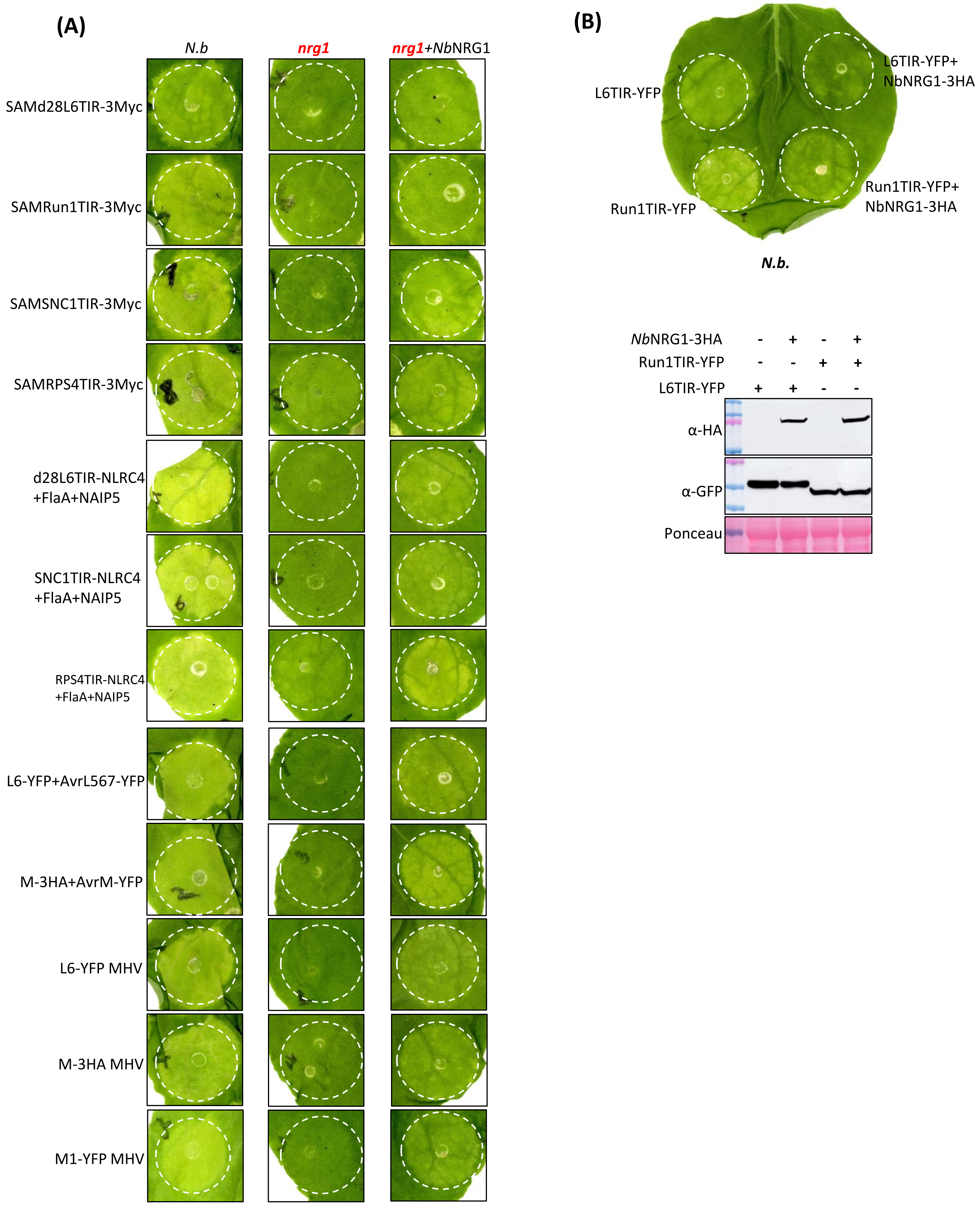


**Supplemental Figure S6. *Nb*NRG1 is required for TIR-mediated cell death in *N. benthamiana*.** (A). Cell death recovery in *nrg1* triple mutant line by Agrobacterium-mediated transient expression of *NbNRG1*. Indicated proteins that fused with YFP tag were expressed alone or together with *Nb*NRG1 that fused with 3xHA tag. Photos were taken at 5 dpi. (B). *Nb*NRG1 inhibited autoactive TIRs induced cell death in WT plants (Infiltration of *Nb*NRG1; OD600 ≥ 0.5). Photos were taken at 5 dpi. Western blot shows the accumulation of TIR proteins with and without *Nb*NRG1.

### Supplemental Figure 7

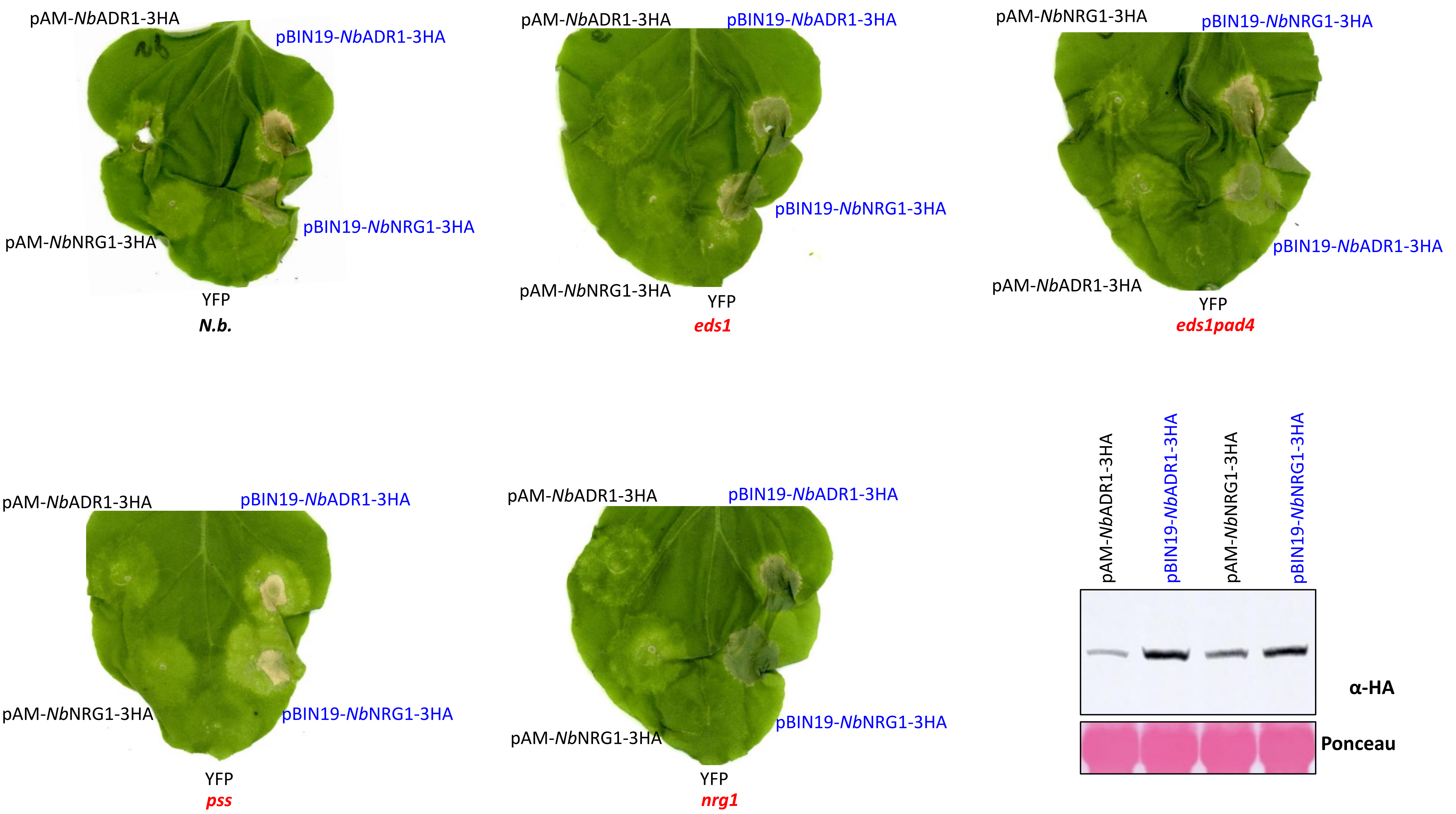


**Supplemental Figure S7. Overexpression of *Nb*NRG1 and *Nb*ADR1 fused to 3xHA tag induce cell death in *N. benthamiana.*** Autoactivity of *Nb*NRG1 and *Nb*ADR1 expressed from pAM-PAT-35s-GWY-3xHA or pBIN19-35s-GWY-3xHA vectors tested in WT and mutant *N. benthamiana* plants (OD600 = 2.0). Western blot showed the accumulation of *Nb*NRG1, *Nb*ADR1 and *Nb*EDS1 proteins from each vector.

### Supplemental Figure 8

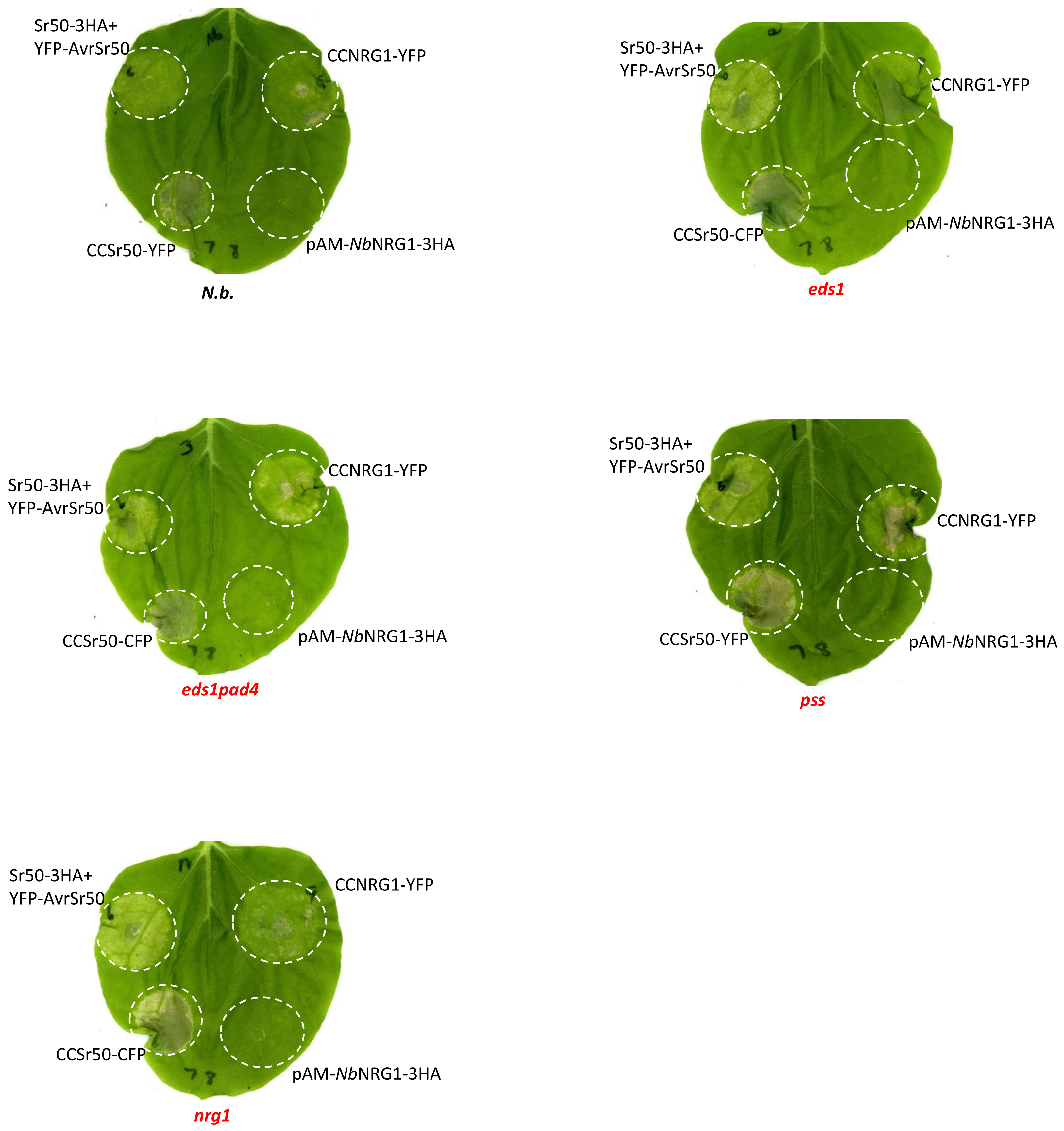


**Supplemental Figure S8. CC-NLRs and autoactive CC domains induce cell death in *N. benthamiana* mutant lines.** Cell death was tested in *N. benthamiana* WT, *eds1*, *eds1pad4*, *pss* and *nrg1* mutant lines. Full length Sr50 (OD600 = 0.1) was co-infiltrated with YFP-AvrSr50 (OD600 = 0.5). CCSr50-YFP (OD600 = 0.5) is the autoactive CC domain from Sr50. CCNRG1-YFP is the autoactive CC domain from *Nb*NRG1. pAM-*Nb*NRG1-3xHA (OD600= 0.5) is not autoactive due to low expression from the construct.

#### Supplemental Figure 9

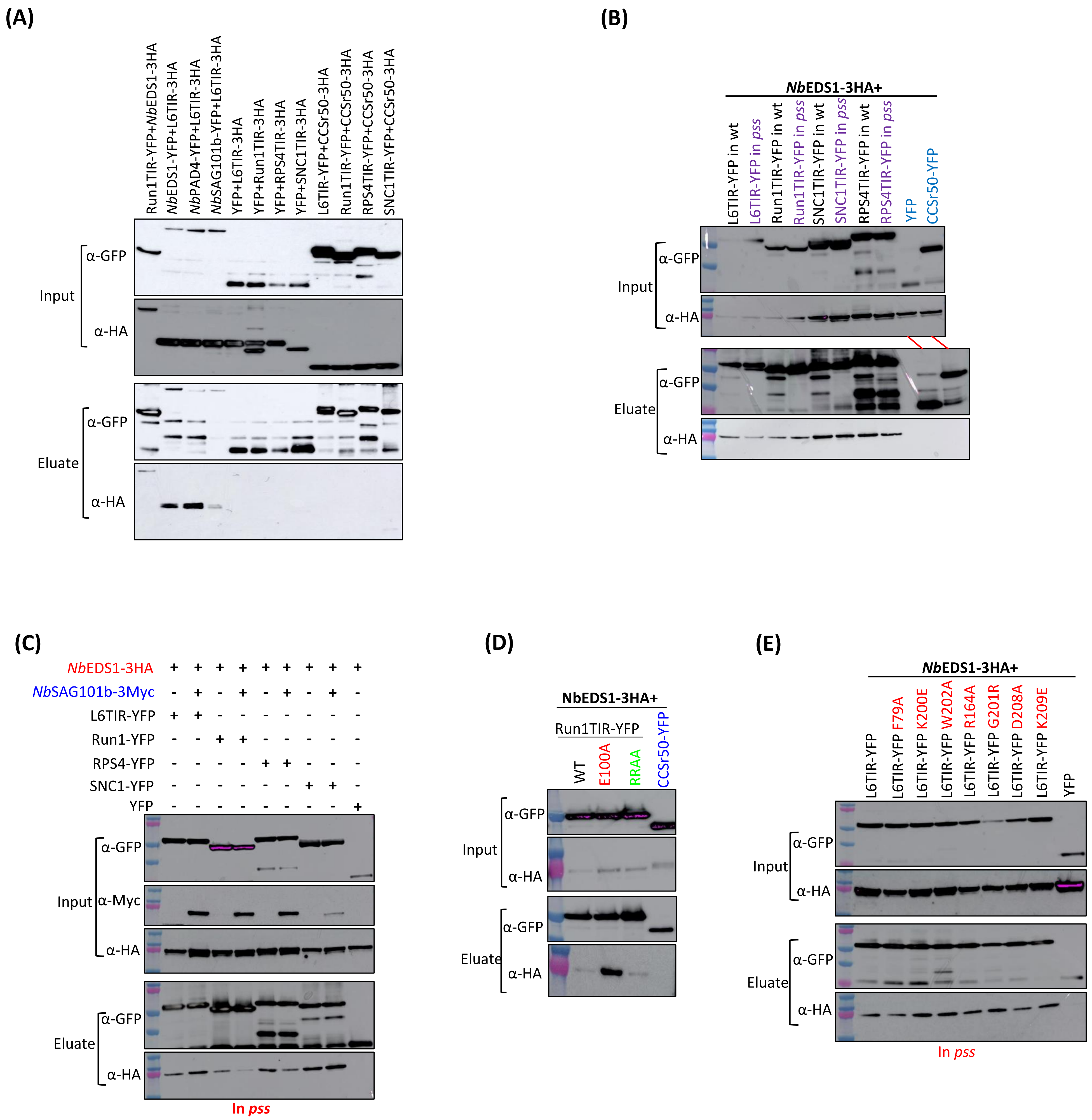


**Supplemental Figure S9. Plant TIRs interact with *Nb*EDS1.** (A). CoIP showed L6TIR-3xHA interacts with *Nb*EDS1-YFP, *Nb*PAD4-YFP and *Nb*SAG101b-YFP. TIRs-3xHA in combination with YFP alone and TIRs-YFP in combination with CC domain of Sr50 that fused to 3xHA tag were used as negative controls. (B). CoIP of *Nb*EDS1-3xHA with wildtype Run1TIR-YFP, Run1TIR-YFP E100A (NADase catalytic site mutations) and Run1TIR-YFP RRAA (R64AR65A), the combinations were transiently expressed in *N. benthamiana* leaves. (C). CoIP of *Nb*EDS1-3xHA with wildtype L6TIR-YFP and AE/DE interfaces mutations, the combinations were expressed in *pss* mutant line. (D). CoIP of *Nb*EDS1-3xHA with plant TIRs-YFP. The combinations were transiently expressed in both wildtype and *pad4/sag101a/sag101b* mutant plants. (E). Competition CoIP experiments of *Nb*EDS1-3xHA with TIR-YFP proteins with and without *Nb*SAG101b-3Myc.

### Supplemental Figure 10

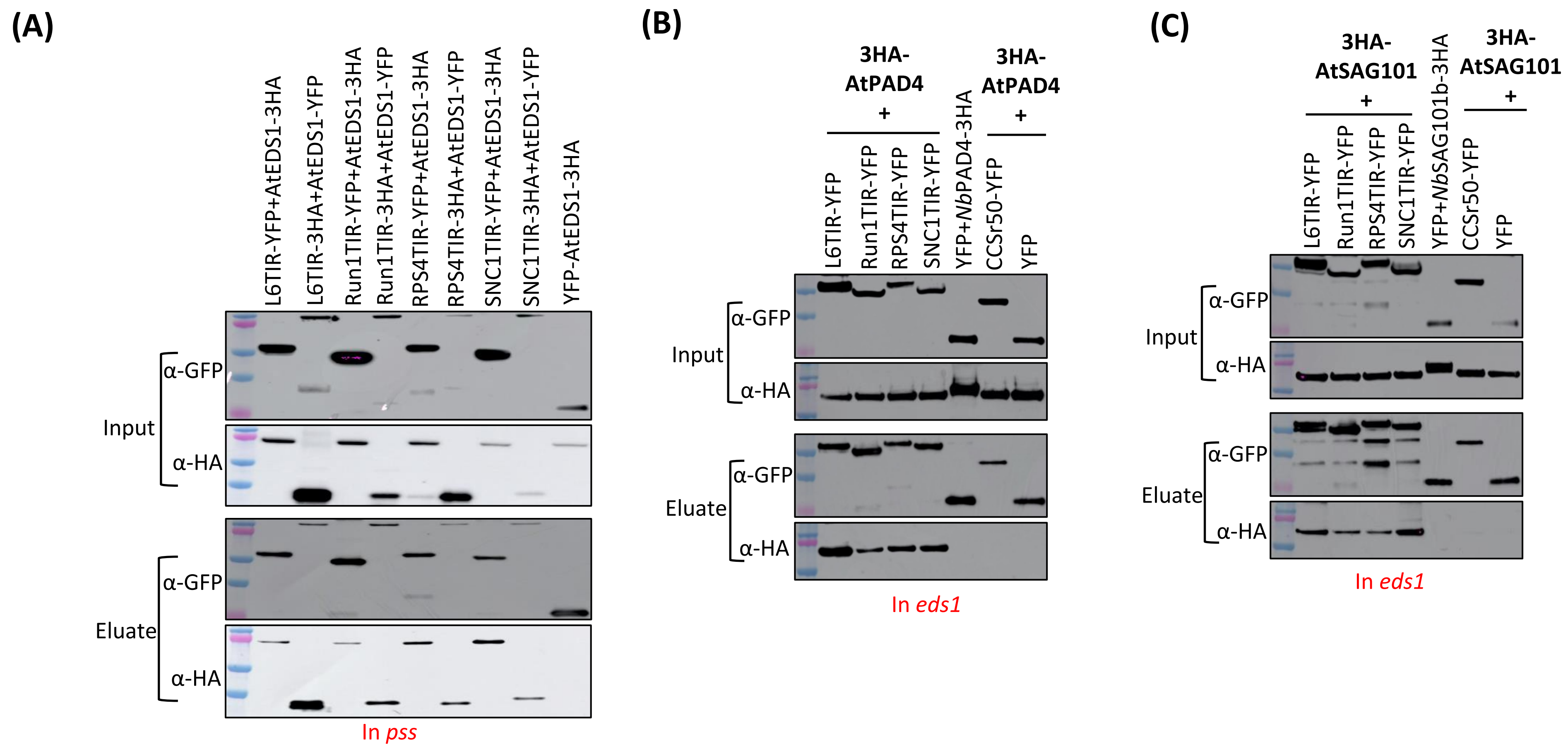


**Supplemental Figure S10. Plant TIRs interact with *Arabidopsis* EDS1, PAD4 and SAG101 in CoIP.** (A) CoIP experiment of *At*EDS1 and different TIRs. The combinations of *At*EDS1-3xHA/-YFP with wildtype TIR-YFP/-3xHA were transiently expressed in *pss* mutant plants. (B) CoIP experiment of *At*PAD4, *At*SAG101 with different plant TIRs in *eds1* plants. *At*PAD4, *At*SAG101 were fused to N-terminal 3xHA tag, TIRs were fused to a YFP tag in the C-terminal of TIR proteins. Leaf samples were collected 1 dpi.

### Supplemental Figure 11

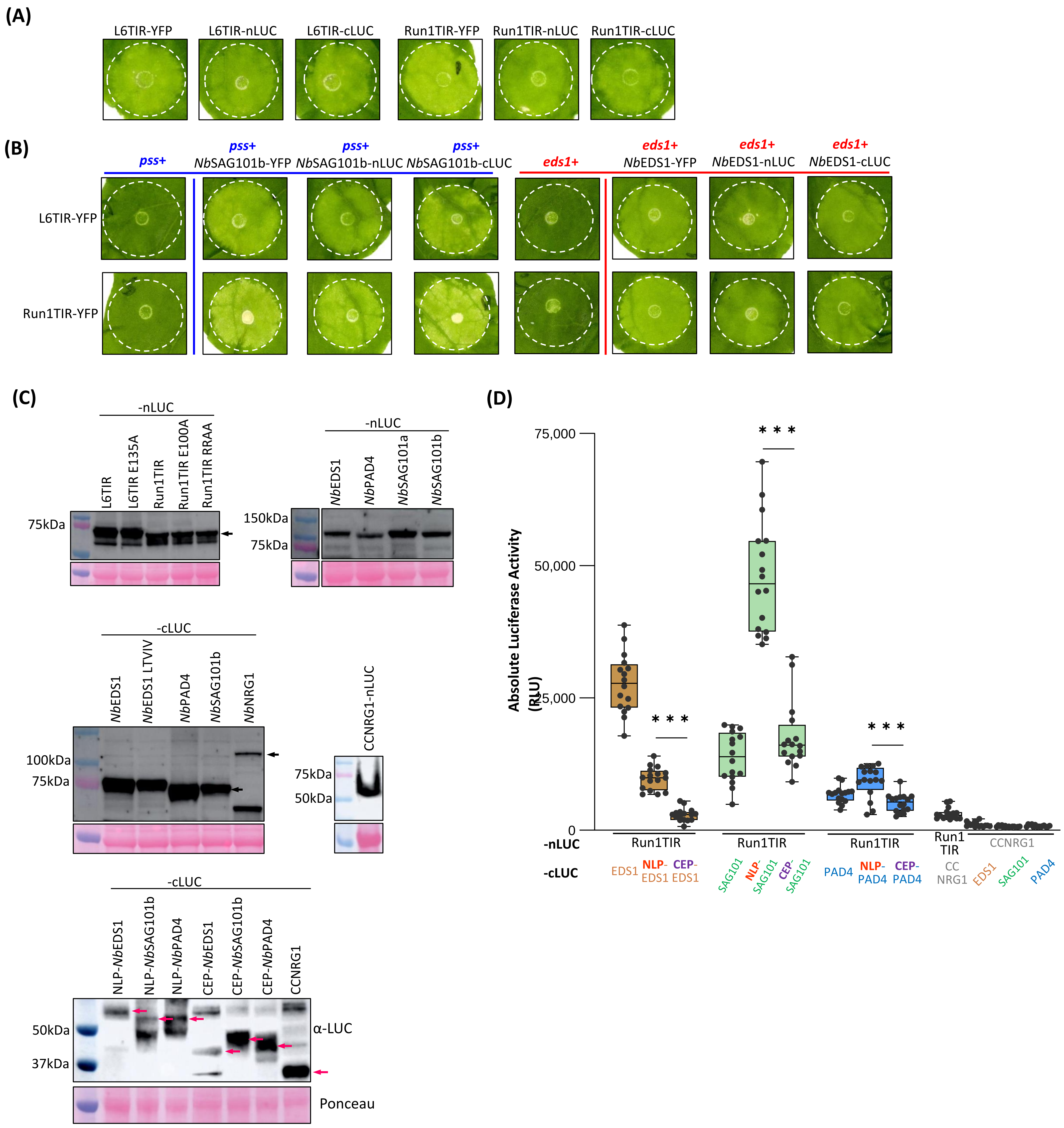


**Supplemental Figure S11. Functional assays of split-luciferase constructs in *N. benthamiana*.** (A). Cell death of L6TIR and Run1TIR fused with cLUC and nLUC in *N. benthamiana*. (B). Cell death complementation assays of L6TIR and Run1TIR fused with YFP tags by *Nb*SAG101b and *Nb*EDS1 fused with cLUC and nLUC in *pss* and *eds1* mutant lines. (C). Protein accumulation from expression of -nLUC and -cLUC fusion constructs**.** All constructs were transiently expressed in *N. benthamiana* plants. Leaf samples were collected 2 dpi. Total proteins were extracted and separated by SDS-PAGE. Western blot membranes were incubated with an anti-luciferase antibody. -cLUC indicates fusion to the C-terminal fragment of luciferase, -nLUC indicates fusion to the N-terminal fragment of luciferase. (D) Split-LUC assays of NbEDS1, NbSAG101b, NbPAD4 sub-domains fused to cLUC with Run1TIR-nLUC. cLUC indicates C-terminal fragment of luciferase, nLUC indicates the N-terminal fragment of luciferase, NLP indicates N-terminal lipase-like domain, CEP indicates C-terminal EP domain. Leaf samples were taken at 2 dpi. The significant difference of luciferase with Run1TIR-nLUC between NLP and CEP domains were labelled with asterisk (***: p<0.001). The combinations of TIR-nLUC+EDS1-cLUC were infiltrated in the *pss* mutant plants, the other combinations were infiltrated in the *eds1* mutant plants.

### Supplemental  Figure 12

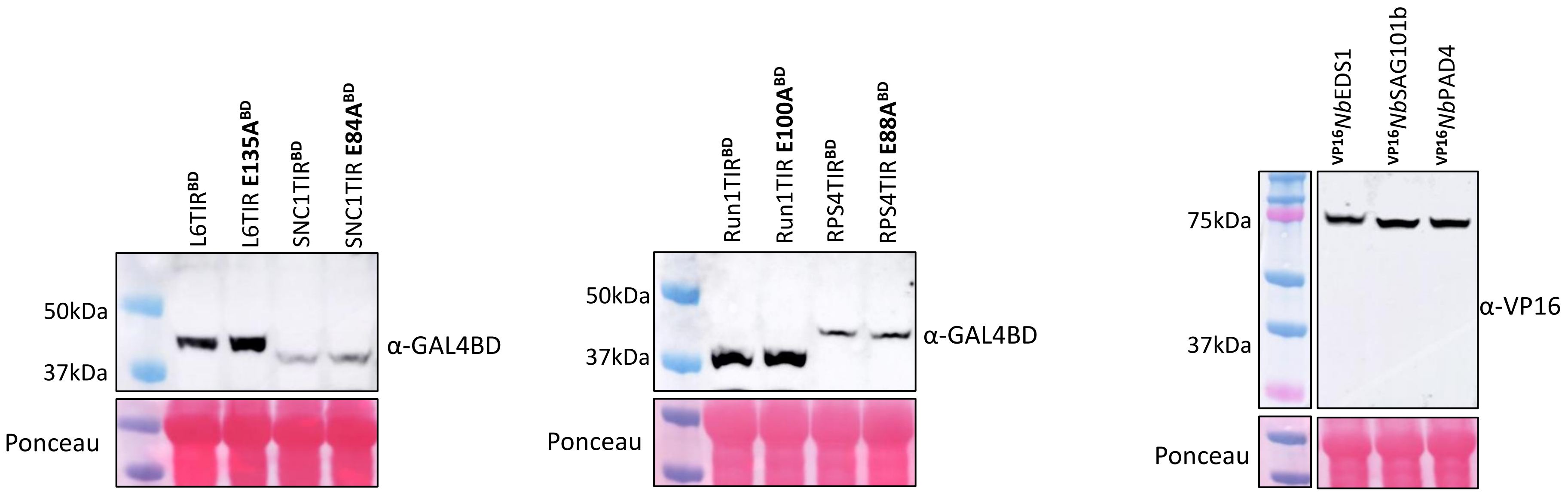


**Supplemental Figure S12. Protein accumulation for expression of plant two-hybrid constructs.** All constructs were transiently expressed in *N. benthamiana* plants, the TIR constructs were infiltrated in *eds1* mutant plants. Leaf samples were collected 2 dpi. Total proteins were extracted and separated by SDS-PAGE. Western blot membranes were incubated with an anti-GAL4BD or anti-VP16 antibodies.

### Supplemental Figure 13

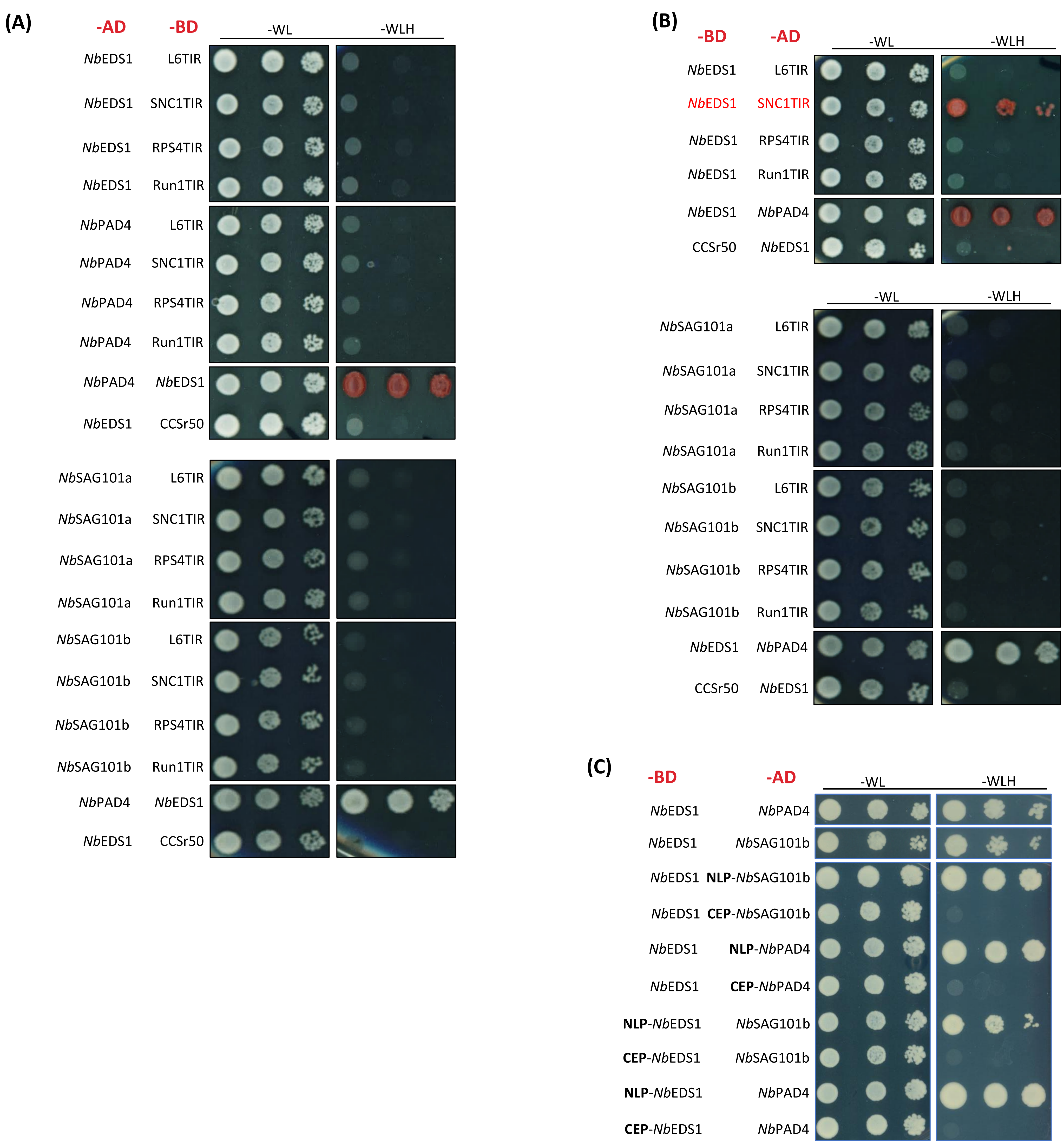


**Supplemental Figure S13. Tests for interaction between plant TIRs and *Nb*EDS1 family proteins by yeast two-hybrid.** (A). Yeast two-hybrid interaction assays between *Nb*EDS1 family proteins fused to the AD domain and plant TIRs fused to the BD domain. (B). Yeast two-hybrid interaction assay between *Nb*EDS1 family proteins fused to the BD domain and plant TIRs fused to the AD domain. *Nb*EDS1-BD+ *Nb*PAD4-AD was used as positive control.C). Heterodimers of full length and truncated *Nb*EDS1 family proteins in Y2H. The full length and truncated *Nb*EDS1 proteins were fused to the BD domain, the full length and truncated *Nb*SAG101b and *Nb*PAD4 proteins were fused to the AD domain. NLP indicates N-terminal lipase-like domain, CEP indicates C-terminal EP domain. Yeast colonies were diluted and grown on media lacking tryptophan and leucine (-WL, growth control), or additionally lacking histidine (-WLH, interaction selection). Pictures were taken after 3 days of growth at 30°C.

### Supplemental Figure 14

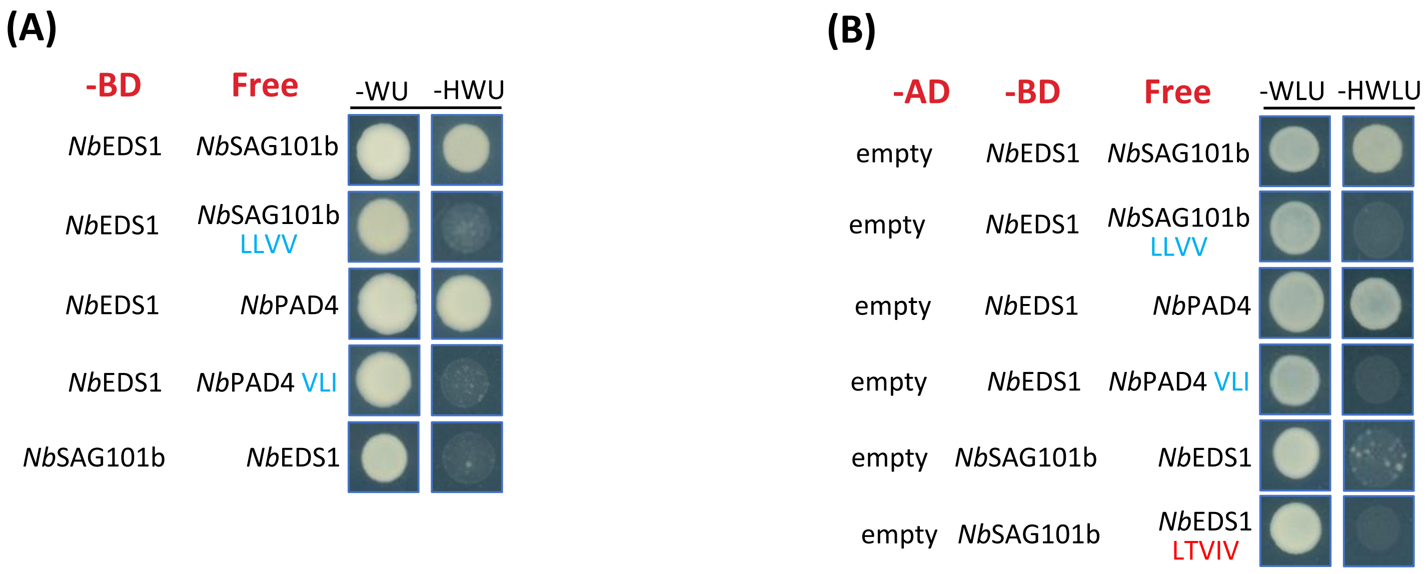


**Supplemental Figure S14. *Nb*EDS1-BD activated yeast HIS3 reporter gene expression when expressed with free *Nb*SAG101b and *Nb*PAD4.** (A). Yeast two-hybrid interaction assay for *Nb*EDS1-BD with free *Nb*SAG101b and *Nb*PAD4 variants and *Nb*SAG101b-BD with free *Nb*EDS1. (B). Yeast three-hybrid assay of AD domain in combination with *Nb*EDS1-BD and *Nb*SAG101b-BD with free *Nb*EDS1, *Nb*SAG101b and *Nb*PAD4 variants. Yeast colonies were diluted and grown on media lacking tryptophan and uracil (-WU, growth control) and media lacking tryptophan, leucine and uracil (-WLU, growth control), or additionally lacking histidine (-HWU, growth control) and (-HWLU, interaction selection). Pictures were taken after 3 days of growth at 30°C.

### Supplemental Figure 15

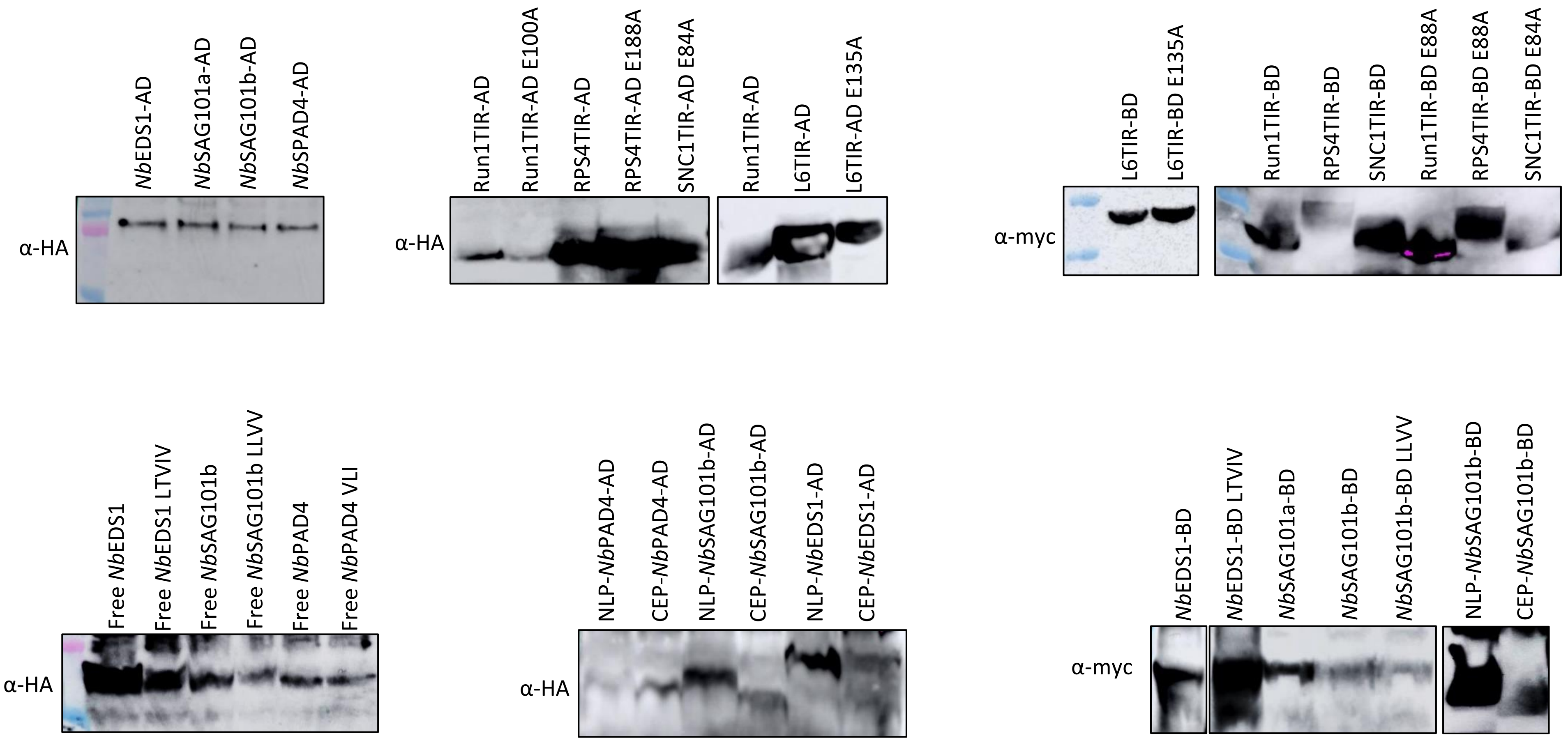


**Supplemental Figure S15. Constructs used in yeast assays have protein expression.** The expression of constructs used in yeast assays were detected by immunoblot analysis. AD fusion and 'free' proteins contain HA tags and were detected using Anti-HA antibody. BD fusion proteins contain a Myc tag and were detected using Anti-Myc antibody.

### Supplemental Figure 16

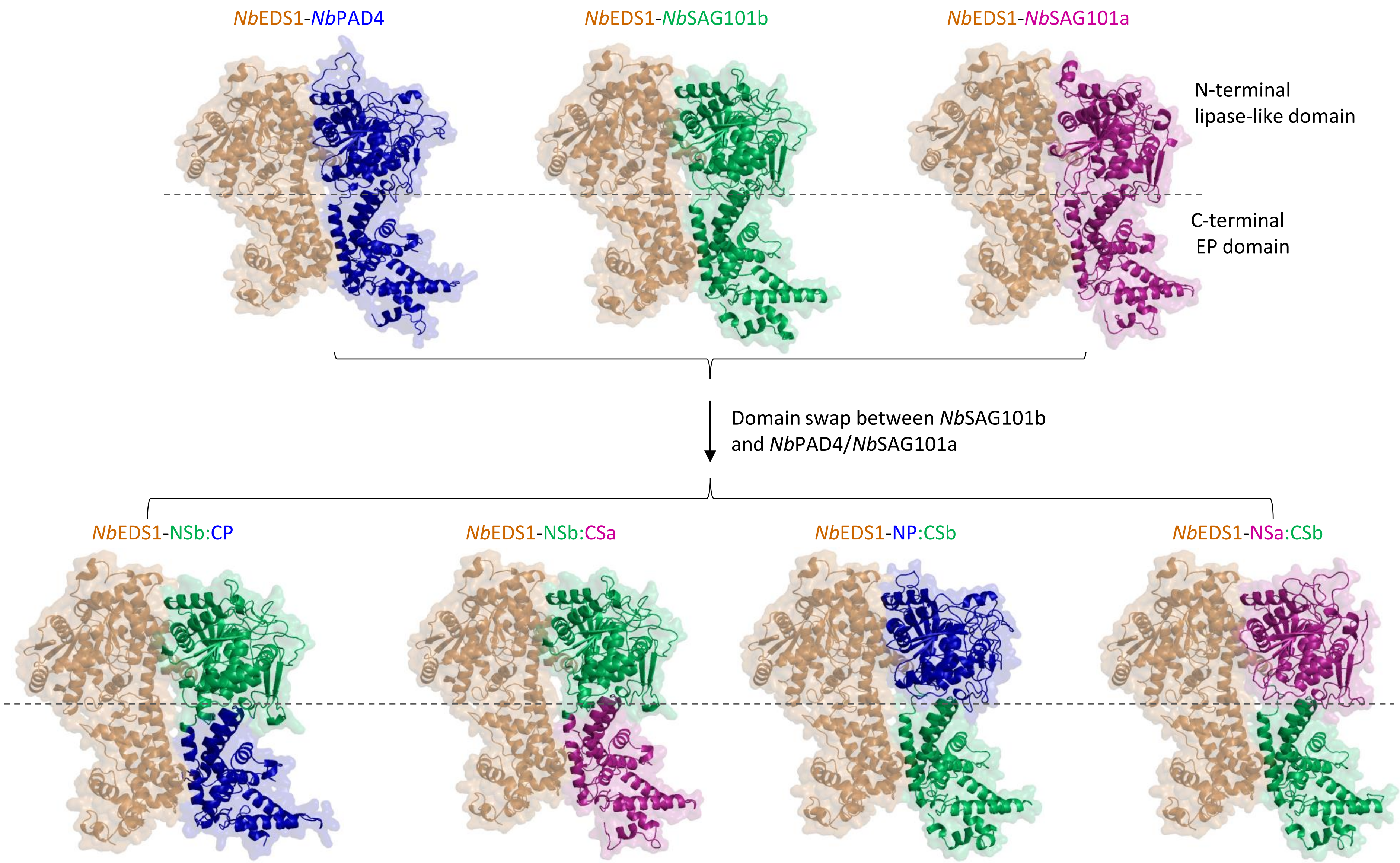


**Supplemental Figure S16. Homology modelling structures of *Nb*EDS1 heterocomplexes with chimeric proteins.** Modelling structures of *Nb*EDS1 heterocomplexes and domain swap chimeras. NSa: N-terminal lipase domain of *Nb*SAG101a; NP: N-terminal lipase domain of *Nb*PAD4; CSb: C-terminal domain of *Nb*SAG101b; CSa: C-terminal domain of *Nb*SAG101a; CP: C-terminal domain of *Nb*PAD4. Structures were modelled on the Arabidopsis *At*EDS1-*At*SAG101 crystal structure (PDB 4nfu) using SWISS-MODEL. Pictures were made using PyMOL (NSb: N-terminal lipase domain of NbSAG101b (aa 1-336); NSa: N-terminal lipase domain of NbSAG101a (aa 1-341); NP: N-terminal lipase domain of NbPAD4 (aa 1-360); CSb: C-terminal domain of NbSAG101b (aa 337-581); CSa: C-terminal domain of NbSAG101a (aa 342-583); CP: C-terminal domain of NbPAD4 (aa 361-609).

### Supplemental Figure 17

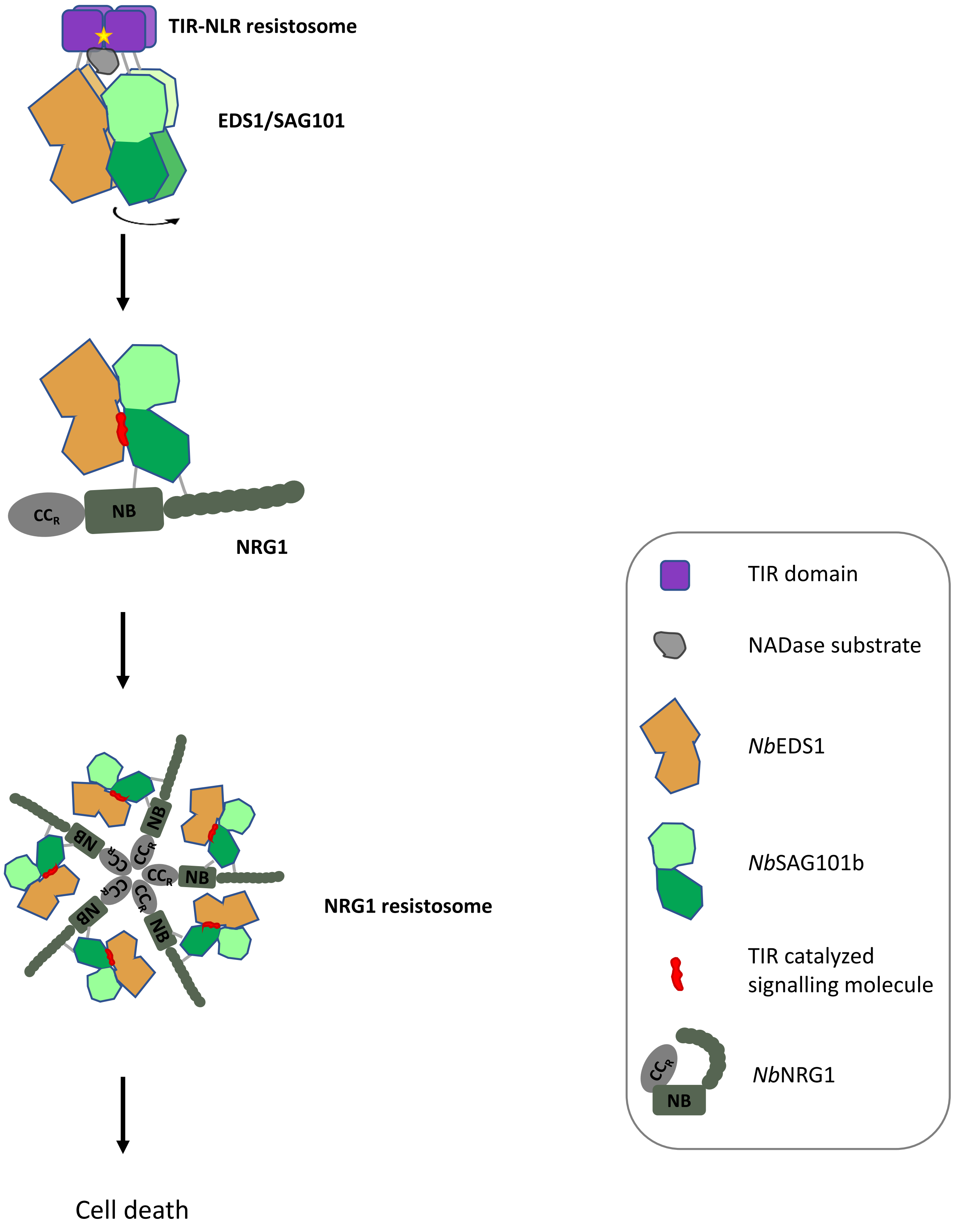


**Supplemental Figure S17. Model for signal transduction in TIR-EDS1/SAG101b/NRG1 pathway.** Proposed model for signal transduction in TIR-*Nb*EDS1/*Nb*SAG101/*Nb*NRG1 pathway. Monomeric TIR domains in inactive TIR-NLRs are capable of interacting with all EDS1 family proteins via their N-terminal lipase-like domains. Upon activation of the TIR-NLR, the active TIR oligomers assemble on the *Nb*EDS1 heterodimers, ensuring that the TIR catalytic function is active in close proximity to the EDS1/SAG101 heterodimer. Once the unstable small signalling molecules are generated they are bound the CEP pocket of the EDS1/SAG101 complex, inducing an allosteric change in the *Nb*SAG101b CEP domain. This allows interaction of the exposed CEP of *Nb*SAG101b with *Nb*NRG1 to activate *Nb*NRG1 leading to formation of the active *Nb*NRG1 resistosome.
